# An IL-2 mutein increases IL-10 and CTLA-4-dependent suppression of dendritic cells by regulatory T cells

**DOI:** 10.1101/2023.12.01.569613

**Authors:** Braxton L. Jamison, Matthew Lawrance, Chun Jing Wang, Hannah A. DeBerg, David M. Sansom, Marc A. Gavin, Lucy S.K. Walker, Daniel J. Campbell

## Abstract

Interleukin-2 (IL-2) variants with increased CD25 dependence that selectively expand Foxp3^+^ regulatory T (T_R_) cells are in clinical trials for treating inflammatory diseases. Using an Fc-fused IL-2 mutein (Fc.IL-2 mutein) we developed that prevents diabetes in non-obese diabetic (NOD) mice, we show that Fc.IL-2 mutein induced an activated T_R_ population with elevated proliferation, a transcriptional program associated with Stat5- and TCR-dependent gene modules, and high IL-10 and CTLA-4 expression. Increased IL-10 signaling limited surface MHC class II upregulation during conventional dendritic cell (cDC) maturation, while increased CTLA-4-dependent transendocytosis led to the transfer of CD80 and CD86 costimulatory ligands from maturing cDCs to T_R_ cells. In NOD mice, Fc.IL-2 mutein treatment promoted the suppression of cDCs in the inflamed pancreas and pancreatic lymph nodes resulting in T cell anergy. Thus, IL-2 mutein-expanded T_R_ cells have enhanced functional properties and restrict cDC function, offering promise for targeted immunotherapy use in autoimmune disease.

## Introduction

Regulatory T (T_R_) cells expressing the transcription factor forkhead box P3 (Foxp3) are required to suppress immune responses against self-antigens and prevent autoimmunity. The cytokine interleukin-2 (IL-2) supports the development, maintenance, and function of Foxp3^+^ T_R_ cells (*1*). There is also an established role of IL-2 in promoting effector and memory CD8^+^ and CD4^+^ T cell responses in infection and anti-tumor immunity (*2, 3*). The opposing functions of IL-2 are explained by varying IL-2 receptor (IL-2R) expression patterns and responsiveness thresholds among cell types. At low concentrations, IL-2 stimulates Foxp3^+^ T_R_ cells constitutively expressing the high-affinity IL-2R (CD25, CD122, and CD132) (*4*). In contrast, memory T cells and natural killer cells expressing the low-affinity IL-2R (CD122 and CD132) are activated at higher concentrations. Furthermore, effector T cells upregulate CD25 after activation, allowing them to compete with T_R_ for limiting amounts of IL-2. Low-dose IL-2 has been explored as a potential treatment for autoimmune diseases due to its ability to boost the expansion of Foxp3^+^ T_R_ cells (*4*). However, the effects on both regulatory and effector T cell populations make balancing the safety and efficacy of low-dose IL-2 challenging, and it has shown inconsistent results in clinical trials (*5–7*), prompting the search for more targeted therapeutic approaches.

There is strong interest in the development of engineered variants of IL-2 (known as IL-2 mutant proteins or muteins) designed to preferentially stimulate Foxp3^+^ T_R_ cells with less off-target specificity. One approach is to decrease the affinity of IL-2 for CD122, thereby increasing CD25 dependence and improving the T_R_ selectivity (*8*). In recent work, we introduced two amino acid substitutions into murine IL-2 that reduce CD122 binding (N103R and V106D, equivalent to the N88 and V91 positions of human IL-2) to create a CD25-biased IL-2 mutein (*9*). To improve pharmacokinetic properties this IL-2 mutein (Fc.Mut24) was fused to an IgG2a Fc domain mutated to reduce FcR binding and effector function. Although Fc.Mut24 is a weaker agonist of IL-2R signaling than an Fc-fused wild-type IL-2 (Fc.WT) *in vitro*, it is significantly better at expanding Foxp3^+^ T_R_ *in vivo* due in part to decreased receptor-mediated clearance that results in an extended biological half-life and sustained IL-2R signaling (*9*). Furthermore, unlike Fc.WT, Fc.Mut24 can be administered at high doses while maintaining T_R_-selectivity and halting disease progression in the non-obese diabetic (NOD) model of Type 1 Diabetes. Several human Fc-fused IL-2 (Fc.IL-2) muteins with similar mutations to Fc.Mut24 are in early-stage clinical development for treating autoimmune disorders (*10*), and phase 1 trials have demonstrated substantial T_R_ expansion and safety (*11*).

Given that clinical trials with T_R_-selective Fc.IL-2 muteins are in progress, it is critical to better understand their mechanism of action (MOA) to optimize therapeutic use. One potential benefit of Fc.IL-2 muteins over other therapies for inflammatory diseases, such as biologics that block proinflammatory cytokines, is their ability to induce durable tolerance by increasing the balance between T_R_ cells and conventional T (T_conv_) cells at the site of tissue inflammation during critical windows of disease development (*9*). An increased T_R_ to effector T cell ratio is associated with anergy induction (*12*), a tolerance mechanism resulting from antigen recognition without sufficient costimulatory signals (*13*). Moreover, T_R_ cells are required for the *in vivo* induction of anergy (*14, 15*). A key feature of T_R_-mediated suppression is the counter-regulation of antigen-presenting cell (APC) function. T_R_ cells are primary producers of the immunosuppressive cytokine interleukin-10 (IL-10), a potent inhibitor of APCs such as macrophages and dendritic cells (DCs) (*16, 17*). Additionally, the checkpoint receptor CTLA-4 is required for T_R_ suppressive activity and functions in a cell-extrinsic manner by capturing the costimulatory ligands CD80 and CD86 that it shares with CD28 in a process termed CTLA-4-dependent transendocytosis (*18–20*). In this process, CD80 and CD86 are transferred from the APC membrane to intracellular compartments in T_R_ and undergo subsequent degradation. Thus, T_R_ cells may promote anergy in autoreactive T_conv_ cells by inhibiting APC maturation via IL-10 and restricting CD28 costimulation via CTLA-4.

Our development of Fc.Mut24 provided a tool to test whether sustained IL-2R signaling influenced T_R_ activation in addition to expansion. We found that Fc.Mut24 treatment induced a novel transcriptional, phenotypic, and functional state in T_R_ which suggested that heightened suppression of APCs may be important for Fc.Mut24-mediated tolerance induction. Using *in vivo* models of DC maturation and examining prediabetic NOD mice, we found that Fc.Mut24 treatment leads to the downregulation of CD80 and CD86 on DCs via CTLA-4-dependent transendocytosis by Foxp3^+^ T_R_ cells, in addition to the downregulation of MHC class II through IL-10 production. The modulation of DC function was associated with T cell anergy and CD73 expression by CD4^+^ and CD8^+^ T_conv_ in NOD mice. Our study links IL-2R signaling in T_R_ with IL-10-mediated suppression of DCs and CTLA-4-dependent transendocytosis, thereby demonstrating how three molecular pathways known to play essential roles in immune regulation converge on the induction of T cell tolerance.

## Results

### Induction of Stat5 and TCR-dependent programs in Foxp3^+^ T_R_ with Fc.Mut24

To investigate how sustained IL-2R signaling with Fc.Mut24 impacted the transcriptional state of T_R_ cells, we treated C57BL/6 (B6) Foxp3-mRFP mice with PBS (vehicle control) or a single 10 µg dose of either Fc.WT or Fc.Mut24. As expected, T_R_ percentages in the spleen significantly increased 3 days after Fc.Mut24 treatment compared to mice given PBS or Fc.WT (Fig 1A). We performed single-cell RNA-seq (scRNA-seq) using the 10x genomics platform on sorted CD4^+^ Foxp3-mRFP^+^ splenic T_R_ cells with four mice per treatment group. After filtering and normalization, 42,698 T_R_ cells were recovered with similar numbers in each group.

**Figure 1.**
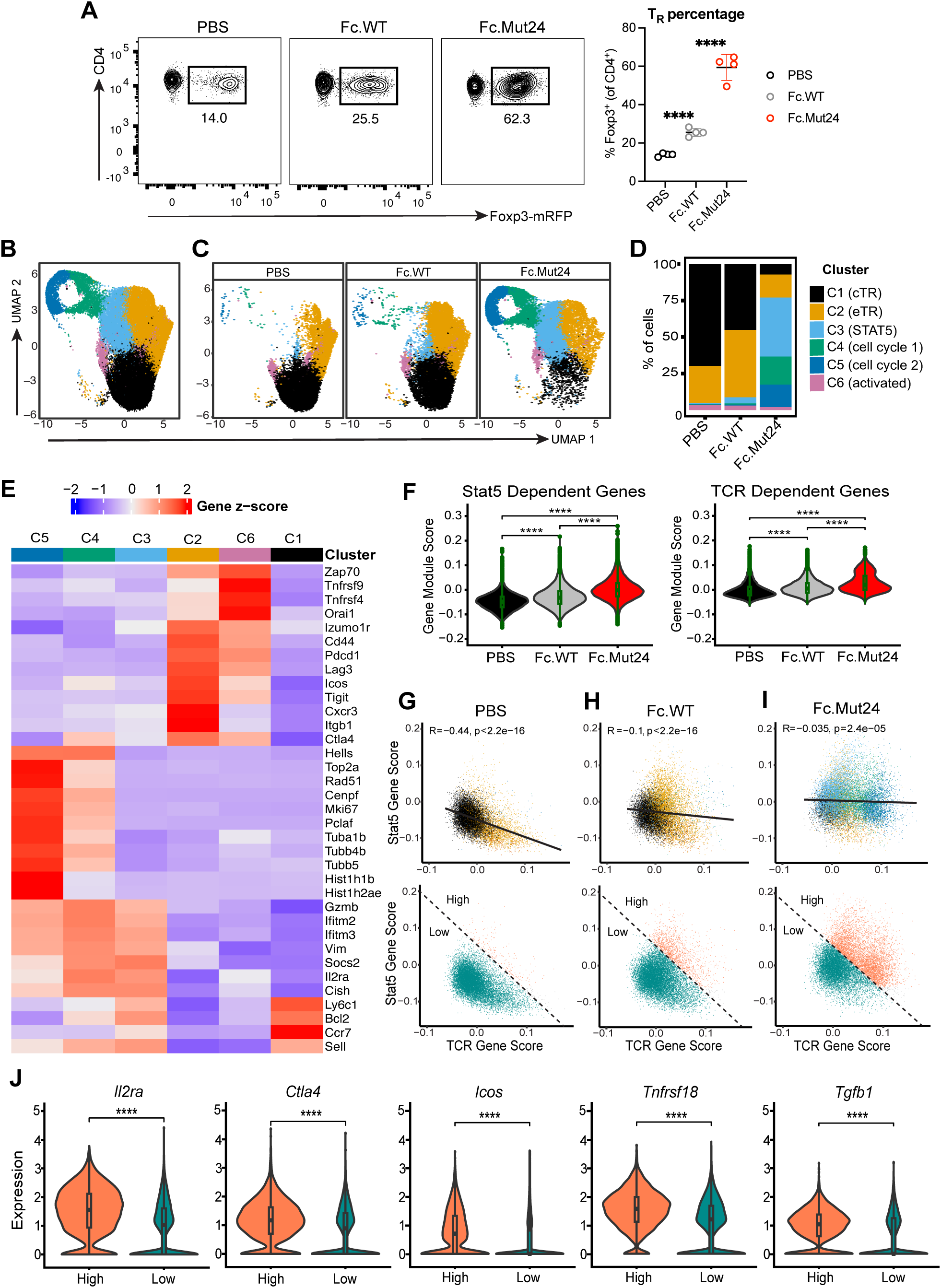
Fc.Mut24 induces a distinct Foxp3^+^ T_R_ cell transcriptional state. Splenocytes from C57BL/6 Foxp3-mRFP treated with PBS, Fc.WT, or Fc.Mut24 were harvested 3 days later and sorted to obtain CD4^+^ Foxp3^+^ T_R_ cells for scRNA-seq. **A.** Percentage of Foxp3^+^ T_R_ from different treatment groups. Representative flow cytometry plots from sorting are shown. Gates were set on live, CD4^+^ cells. Graph shows mean ± SD with individual data points (n = 4); \*\*\*\**P* ≤ 0.0001, two-tailed unpaired t-tests. **B.** UMAP-based clustering of merged scRNA-seq profiles of T_R_ from PBS control, Fc.WT-treated, and Fc.Mut24-treated mice. Individual cells are colored by cluster assignments. A complete list of the differentially expressed genes for each cluster is available in Table S1. **C.** UMAP-based clustering of scRNA-seq profiles of T_R_ by treatment group **D.** Bar graph showing the frequency of T_R_ clusters in each treatment group. **E**. Heat map showing the mean expression (z-score) of representative signature genes (rows) for each cluster (columns). **F.** Violin plots with gene module expression for Stat5-dependent genes (Chinen T et al., *Nat Immunol*, 2016) or TCR-dependent genes (Levine et al., *Nat Immunol*, 2014) in T_R_ cells. **G-I.** Biaxial plots showing expression of Stat5-dependent versus TCR-dependent genes in each treatment group colored by T_R_ cluster (top) or using a high and low cutoff based on co-expression of both modules (bottom). **J.** Violin plots showing normalized expression of select genes associated with T_R_ activation between high and low populations from all treatment groups based on cutoff in G-I (bottom row).

Uniform manifold approximation and projection (UMAP) analysis and unsupervised graphical clustering based on features of all cells organized the Foxp3^+^ T_R_ into six clusters (Fig. 1B-D). Whereas the vast majority of cells from PBS controls or Fc.WT-treated mice were found in Clusters 1 and 2, T_R_ from Fc.Mut24-treated mice were predominantly found in Clusters 2, 3, 4, and 5 (Fig. 1C&D). A small fraction of T_R_ cells from all treatment groups were found in Cluster 6. These clusters were annotated based on differential gene expression (Fig 1E). Genes previously associated with naïve-like central T_R_ (cT_R_) cells such as *Ly6c1, Bcl2, Ccr7*, and *Sell* were highly expressed in Cluster 1, whereas Cluster 2 expressed *Izumo1r*, *Cd44, Pdcd1, Lag3, Icos, Tigit, and Ctla4* that define effector T_R_ (eT_R_) cells (*21, 22*). Clusters 3, 4, and 5 expressed genes induced by Stat5 activation, including *Gzmb*, *Ifitm2, Ifitm3, Vim, Socs2, Il2ra, and Cish* (*23–25*). Based on the expression of genes controlling cell proliferation and DNA replication or repair, such as *Hells, Rad51, Cenpf, Mki67, and Pclaf,* Clusters 4 and 5 represent actively dividing cells. Genes that play a role in chromatin remodeling or microtubule assembly such as *Top2a, Hist1h1b, Hist1h2ae, Tuba1b, Tubb4b, and Tubb55* were highly expressed in Cluster 5 distinguishing these cells from Cluster 4. Cluster 6 highly expressed genes upregulated by T cell activation, such as *Zap70*, *Tnfrsf9*, *Tnfrsf4*, and *Orai1*.

The cluster annotations were further supported by examining differential transcription factor (TF) expression (Fig S1A), which showed enrichment of *Lef1*, *Tcf7*, and *Bach2* associated with T cell stemness in Cluster 1 (*26, 27*). Cluster 2 expressed TFs associated with T_R_ effector function or stability such as *Hif1a*, *Batf*, *Maf*, and *Ikzf2* (*28–31*), and Cluster 3 had the highest expression of Foxp3, a known target gene of Stat5 in T_R_ cells (*32*). TFs involved in cell cycle progression, such as *E2f1, E2f2, Mybl2, and Tfdp1* were expressed in Clusters 4 and 5, while *Tox*, *Nr4a1*, *Nr4a3*, and *Egr2*, which are associated with NFAT activation downstream of antigen receptor signaling were expressed in Cluster 6 (*33–35*).

In addition to IL-2R signaling that supports the survival of cT_R_ cells (*21*), the abundance and proliferation of eT_R_ cells are controlled by signaling through the T cell receptor (TCR) (*36*). Surprisingly, analysis of genes regulated in T_R_ cells by either Stat5 activation (*37*), or TCR signaling (*36*), showed that Fc.Mut24 treatment upregulated both of these transcriptional programs relative to PBS or Fc.WT (Fig. 1F). In T_R_ from PBS controls, there was an inverse correlation between the expression of Stat5- and TCR-dependent programs, and these gene sets were associated with cT_R_ in Cluster 1 and eT_R_ in Cluster 2, respectively (Fig. 1G). In contrast, T_R_ from Fc.Mut24-treated and to a lesser extent Fc.WT-treated mice co-expressed the Stat5- and TCR-dependent programs, and in response to Fc.Mut24 this was associated with Clusters 3, 4, and 5 (Fig. 1H&I). Cells with high expression of both Stat5- and TCR-dependent programs had increased expression of genes associated with highly functional T_R_, including *Il2ra*, *Ctla4*, *Icos, Tnfrsf18, and Tgfb1* (Fig. 1G-J). Despite the prevalence of a TCR-dependent gene signature in T_R_ from Fc.Mut24-treated mice, single-cell TCR sequencing showed that Fc.Mut24-expanded T_R_ maintained a diverse TCR repertoire that was comparable to PBS controls or Fc.WT-treated mice, and had limited clonal expansion (Fig. S1B&C). These data indicate that expansion of the T_R_ population following Fc.Mut24 treatment is characterized by co-expression of Stat5- and TCR-dependent gene signatures, which is elevated compared to Fc.WT treatment, and associated with activation.

### Unique phenotypic features of Foxp3^+^ T_R_ are induced with Fc.Mut24

Given the striking transcriptional changes induced by Fc.Mut24 treatment, we further analyzed its role in modulating T_R_ phenotype and function. We injected B6 mice with 10 µg of Fc.Mut24 and examined splenic T_R_ cells 3 days later. We observed an ∼8-fold expansion in T_R_ cell number in Fc.Mut24-treated mice compared to mice given PBS (Fig. 2A), and nearly all T_R_ cells were Ki-67^+^ after treatment, indicative of sustained proliferation (Fig. 2A). Fc.Mut24 treatment increased the expression of CD44 (Fig. 2A), a marker associated with T cell activation, previously shown to identify proliferating T_R_ cells in the steady state (*38*). Expression of the hallmark T_R_ identity markers CD25, CTLA-4, and ICOS were also increased (Fig. 2A). In contrast, expression of the coinhibitory receptor PD-1, which antagonizes T_R_ function and is downregulated by Stat5 activation (*39, 40*), did not increase (Fig. 2A).

**Figure 2.**
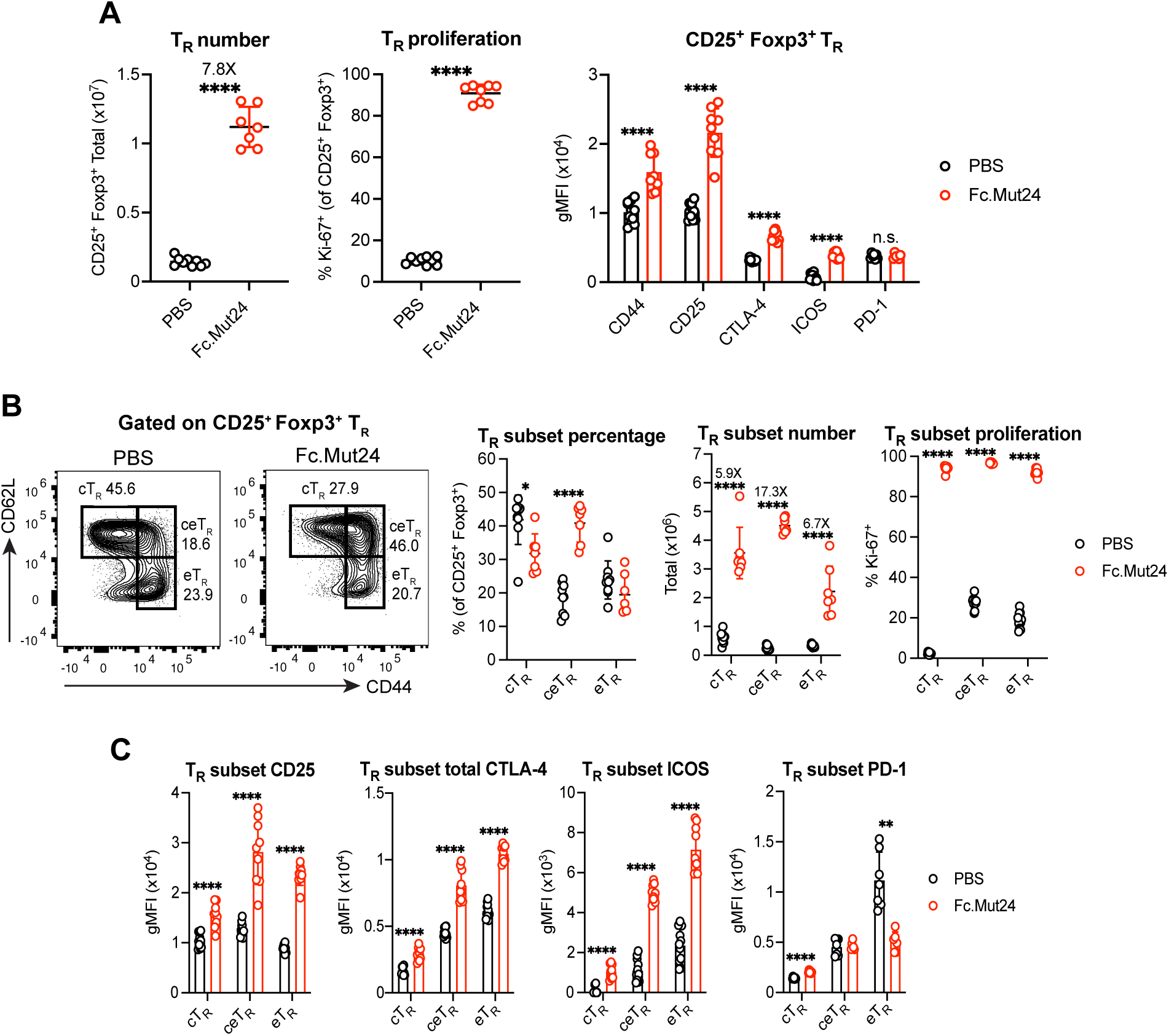
Fc.Mut24 activates different Foxp3^+^ T_R_ cell subsets. Splenocytes from C57BL/6 mice treated with PBS or Fc.Mut24 were harvested 3 days later and analyzed by flow cytometry. **A.** Number and proliferation (Ki-67) of CD25^+^ Foxp3^+^ T_R_ and the expression of indicated T_R_ activation markers. Gates were set on live, CD4^+^ cells. **B.** Percentage, number, and proliferation of CD44^lo^ CD62L^hi^ central T_R_ (cT_R_), CD44^hi^ CD62L^hi^ central effector T_R_ (ceT_R_), and CD44^hi^ CD62L^lo^ effector T_R_ (eT_R_) in each treatment group. Gates are indicated on representative flow cytometry plots. **C.** Expression of indicated T_R_ activation markers on T_R_ subsets. **A-C.** All graphs show mean ± SD with individual data points (n = 6 to 11); \**P* ≤ 0.05, \*\**P* ≤ 0.01, \*\*\*\**P* ≤ 0.0001, two-tailed unpaired t-tests, or multiple unpaired *t*-tests. n.s., not significant. Data are representative of 2-3 independent experiments.

We analyzed CD44^lo^ CD62L^hi^ cT_R_ and CD44^hi^ CD62L^lo^ eT_R_ populations since cT_R_ cells in secondary lymphoid tissues are more dependent on IL-2R signaling during homeostasis (*21, 41*). There was close to an equal expansion of cT_R_ (6-fold) and eT_R_ (7-fold) in response to Fc.Mut24 treatment (Fig. 2B). However, a population of CD44^hi^ CD62L^hi^ T_R_, which we called central effector (ceT_R_) cells, had the largest increase in both percentage and cell number (17-fold) in Fc.Mut24-treated mice (Fig. 2B). Fc.Mut24 treatment upregulated CD25 on both cT_R_ and eT_R_, but similar to the expansion of cell number, ceT_R_ had the highest overall CD25 expression (Fig. 2B). The expression of CTLA-4 and ICOS was increased on all T_R_ subsets after Fc.Mut24 treatment, with the highest levels of these markers observed on eT_R_ followed by ceT_R_ (Fig. 2C). We also found that although Fc.Mut24 treatment had no effect on PD-1 when total T_R_ cells were analyzed, PD-1 expression was specifically reduced in eT_R_ (Fig. 2C), which have the highest baseline PD-1 levels due to strong TCR signaling (*42*).

### CTLA-4 cycling by Foxp3^+^ T_R_ is enhanced with Fc.Mut24

Although CTLA-4 is constitutively expressed by eT_R_, it is largely restricted to intracellular vesicles due to rapid internalization from the plasma membrane via clathrin-mediated endocytosis (*43*). Following TCR stimulation, the intracellular pool of CTLA-4 is mobilized and rapidly cycles from these vesicles to the cell surface, allowing CTLA-4 to bind and strip CD80 and CD86 from APCs via transendocytosis (*19, 20*). Therefore, to follow up our findings that total CTLA-4 expression increased after Fc.Mut24 treatment (Fig. 2A), we measured surface CTLA-4 by staining live cells at 4°C for 30 minutes and stained for the cycling pool of functional CTLA-4 molecules by labeling for 2 hours at 37°C. We found that Fc.Mut24 treatment dramatically increased both surface and cycling CTLA-4 in Foxp3^+^ T_R_ cells but not in Foxp3^-^ conventional T (T_conv_) cells with ∼17% of T_R_ cycling CTLA-4 after treatment (Fig. 3A). Additionally, analysis of cT_R_, ceT_R_, and eT_R_ showed significantly increased CTLA-4 surface expression and cycling in all T_R_ subsets after Fc.Mut24 treatment (Fig. 3B). Consistent with these results, the expression of *Trat1* (TRIM), *Lax1* (LAX), and *Rab8* that form a multimeric complex regulating post-Golgi trafficking of CTLA-4 to the cell surface were upregulated in T_R_ from Fc.Mut24-treated mice (Fig. S2A) (*44*). Fc.Mut24 treatment also increased the expression of the guanine nucleotide exchange factor *Def6* and the small GTPase Rab11 (*Rab11a* and *Rab11b*), which interact to control endosomal recycling of CTLA-4 to the cell surface (Fig. S2B) (*45, 46*).

**Figure 3.**
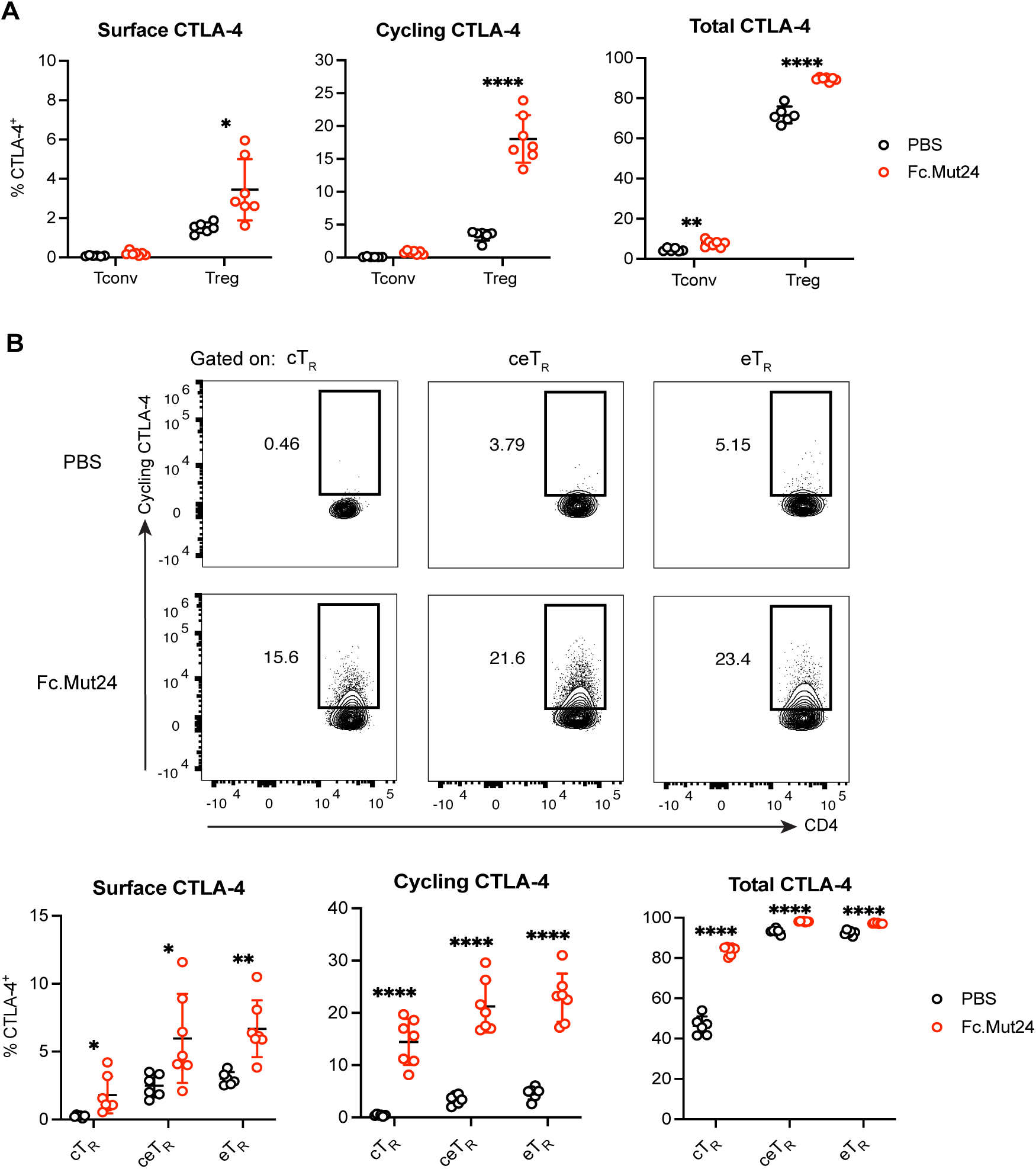
Fc.Mut24 increases CTLA-4 cycling by different Foxp3^+^ T_R_ cell subsets. Splenocytes from C57BL/6 mice treated with PBS or Fc.Mut24 were harvested 3 days later and stained for CTLA-4 expression at 4°C (Surface), at 37°C for 2 h (Cycling), or on fixed and permeabilized cells (Total) followed by analysis by flow cytometry. **A.** Percentage of CTLA-4^+^ cells between different treatment groups. Gates were set on live, CD4^+^, Foxp3^+^ T_R_ or Foxp3^-^ T_conv_ cells. **B.** Percentage of cycling CTLA-4^+^ cells between CD44^lo^ CD62L^hi^ central T_R_ (cT_R_), CD44^hi^ CD62L^hi^ central effector T_R_ (ceT_R_), and CD44^hi^ CD62L^lo^ effector T_R_ (eT_R_). Representative flow cytometry plots for Cycling CTLA-4 are shown. **A&B.** All graphs show mean ± SD with individual data points (n = 6 to 9); \**P* ≤ 0.05, \*\**P* ≤ 0.01, \*\*\*\**P* ≤ 0.0001, multiple unpaired *t*-tests. Data are representative of 2 independent experiments.

### CTLA-4-dependent capture of costimulatory ligands by Foxp3^+^ T_R_ is enhanced with Fc.Mut24

To determine if the increased CTLA-4 cycling in T_R_ cells resulted in increased CTLA-4-dependent transendocytosis of costimulatory ligands, we isolated CD4^+^ T cells from PBS controls or Fc.Mut24-treated mice, co-cultured them with NIH/3T3 fibroblasts expressing GFP-tagged CD86 for 2 hours, and analyzed ligand capture by flow cytometry (Fig. 4A). In accordance with the CTLA-4 cycling results (Fig. 3A), almost no T_conv_ cells from either PBS controls or Fc.Mut24-treated mice captured the CD86-GFP ligand, and only a low level of CD86-GFP capture (<5% of cells) was observed in T_R_ from PBS controls (Fig. 4B). However, this capture significantly increased to ∼17% of T_R_ after Fc.Mut24 treatment (Fig. 4B). Addition of the anti-CTLA-4 blocking antibody clone UC10-4F10-11 (4F10) reduced CD86-GFP capture, and internalization of captured ligand was confirmed by the accumulation of CD86-GFP in the presence of the lysosomal inhibitor Bafilomycin A1 (BafA) (Fig. 4C). In line with a previous report (*20*), we found that eT_R_ exhibited the highest level of CTLA-4 transendocytosis in PBS controls (Fig. S3A&B). However, after Fc.Mut24 treatment ceT_R_ cells captured CD86-GFP ligand as efficiently as eT_R_, and treatment also significantly increased ligand capture by cT_R_ (Fig. S3A&B).

**Figure 4.**
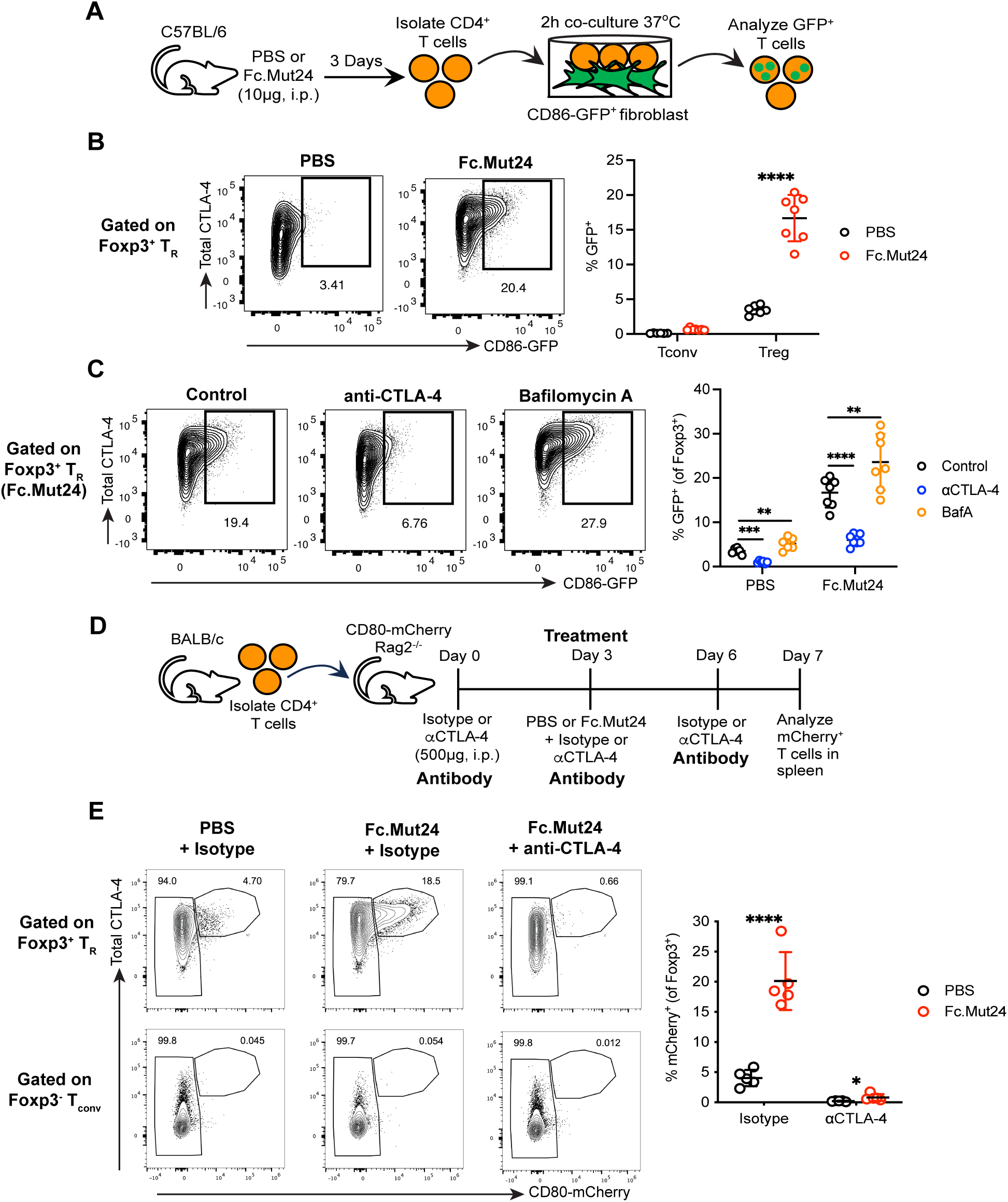
Fc.Mut24 increases CTLA-4-dependent transendocytosis of CD80 and CD86 by Foxp3^+^ T_R_ cells. **A.** Experimental design schematic for *ex vivo* CTLA-4 transendocytosis assay. Gates were set on live, CD4^+^, Foxp3^+^ T_R_ or Foxp3^-^ T_conv_ cells. **B.** Capture of CD86-GFP by Foxp3^+^ T_R_ and Foxp3^-^ T_conv_ between different treatment groups. Representative flow cytometry plots for T_R_ are shown. Graph shows mean ± SD with individual data points (n = 7); \*\*\*\**P* ≤ 0.0001, multiple unpaired *t*-tests. Data are representative of 2-3 independent experiments. **C.** Capture of CD86-GFP by Foxp3^+^ T_R_ between different co-culture conditions. Representative flow cytometry plots from Fc.Mut24-treated mice are shown. Graph shows mean ± SD with individual data points (n = 7); \*\**P* ≤ 0.01, \*\*\**P* ≤ 0.001, \*\*\*\**P* ≤ 0.0001, multiple paired *t*-tests. Data are representative of 2-3 independent experiments. **D.** Experimental design schematic for *in vivo* CTLA-4 transendocytosis assay. Gates were set on CD4^+^, Foxp3^+^ T_R_ or Foxp3^-^ T_conv_. **E.** Capture of CD80-mCherry by Foxp3^+^ T_R_ and Foxp3^-^ T_conv_ between different treatment groups. Representative flow cytometry plots are shown. The graph shows Foxp3^+^ T_R_ data, and mean ± SD with individual data points (n = 5 per group); \**P* ≤ 0.05, \*\*\*\**P* ≤ 0.0001, multiple unpaired *t*-tests. Data are representative of 3 independent experiments.

To show that Fc.Mut24 also enhanced CTLA-4-dependent transendocytosis by T_R_ *in vivo*, we set up a model in which CD4^+^ T cells were adoptively transferred into CD80-mCherry.Rag2^-/-^ mice followed by Fc.Mut24 treatment (Fig. 4D). The CD80-mCherry fusion gene is knocked into the endogenous locus in these recipient animals, allowing for tracking CD80 transfer from antigen-presenting cells (APCs) to T cells (Wang et al., manuscript in preparation). In addition to Fc.Mut24 treatment, recipients received 3 injections of isotype or anti-CTLA-4 blocking antibody (4F10) to confirm the role of CTLA-4 in ligand capture (Fig. 4D). Similar to our *ex vivo* CTLA-4 transendocytosis assay, <5% of Foxp3^+^ T_R_ from PBS controls captured CD80-mCherry ligand *in vivo*, while virtually no T_conv_ cells did (Fig. 4E). However, Fc.Mut24 treatment significantly increased the percentage of CD80-mCherry^+^ T_R_ to ∼20%, with no changes observed for T_conv_, and administration of 4F10 almost completely blocked CD80-mCherry ligand capture by T_R_ (Fig. 4E). Thus, in addition to stimulating T_R_ expansion, Fc.Mut24 enhances T_R_ function by increasing CTLA-4 cycling and transendocytosis of costimulatory ligands by multiple subsets of Foxp3^+^ T_R_ cells.

### Upregulation of costimulatory ligand and MHC class II protein expression during cDC maturation is suppressed with Fc.Mut24

Given the elevated function of CTLA-4 in T_R_ induced by Fc.Mut24 treatment, we examined how this in turn impacted the maturation of different APC populations following inflammatory stimulation. In secondary lymphoid organs, two primary cDC subsets have been described. Development of type 1 conventional DCs (cDC1s) depends on the transcription factors Batf3 and Irf8 (*47*), and cDC1s are essential for cross-presentation and priming of CD8^+^ T cells as well as Th1 cells (*48*). In contrast, the development of type 2 conventional DCs (cDC2s) is Irf4-dependent, and cDC2s prime Th17 and Th2 responses (*49*). To determine the impact of Fc.Mut24 treatment on cDC1 and cDC2 *in vivo*, we treated mice with PBS or Fc.Mut24, and 3 days later some mice received a low dose of Lipopolysaccharide (LPS) intraperitoneally to promote cDC maturation in the spleen (Fig. 5A). Interestingly, whereas Fc.Mut24 treatment had no impact on 33D1^+^ cDC2 frequency in the spleen, both the percentage and number of splenic CD8α^+^ cDC1s were increased after treatment (Fig. S4A-B). However, this expansion was lost in the presence of LPS, where Fc.Mut24 treatment was associated with a decreased abundance of cDC1s.

**Figure 5.**
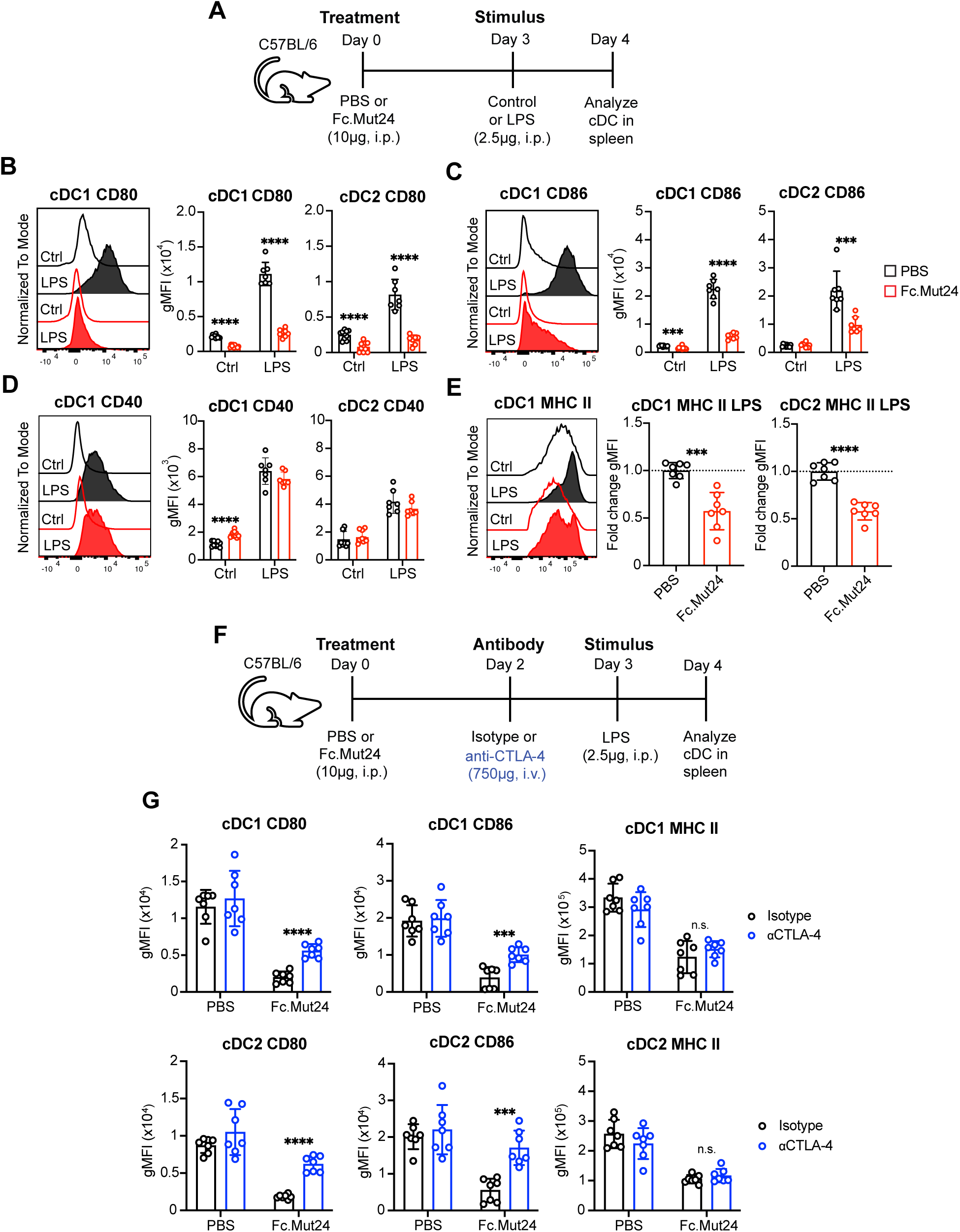
Fc.Mut24 promotes CTLA-4-dependent transendocytosis of CD80 and CD86 during dendritic cell maturation. **A.** Experimental design schematic for the *in vivo* dendritic cell (DC) maturation model. See Fig. S4A for the gating strategy. **B-E.** Expression of indicated DC maturation markers. Representative histograms are shown. Graphs show mean ± SD with individual data points (n = 7 to 10); \*\*\**P* ≤ 0.001, \*\*\*\**P* ≤ 0.0001, multiple unpaired *t*-tests. Ctrl, Control. For MHC class II expression, data are shown as fold-change gMFI over the mean of PBS control (dotted line). Data are representative of 2-3 independent experiments. **F.** Experimental design schematic for the anti-CTLA-4 study. **G.** Expression of indicated DC maturation markers between isotype- and anti-CTLA-4 antibody-treated mice. Graphs show mean ± SD with individual data points (n = 7); \*\*\**P* ≤ 0.001, \*\*\*\**P* ≤ 0.0001, multiple unpaired *t*-tests. n.s., not significant. Data are representative of 2-3 independent experiments.

As expected, LPS stimulation in PBS controls led to classical signs of DC maturation, such as increased surface expression of costimulatory ligands (CD80 and CD86), costimulatory receptors (CD40), and antigen-presentation molecules (MHC class II) on cDC1s and cDC2s (Fig. 5B-E). However, in the context of prior Fc.Mut24 treatment, the LPS-mediated upregulation of CD80 and CD86 on both cDC1s and cDC2s was almost completely blocked (Fig. 5B&C), whereas there was no effect on upregulation of CD40 (Fig. 5D). LPS-mediated upregulation of MHC class II surface expression was partially reduced in both cDC1s and cDC2s in Fc.Mut24-treated mice (Fig. 5E), but not to the same extent as what was observed for CD80 and CD86. In the absence of LPS, the effects of Fc.Mut24 treatment were less dramatic but included increased expression of CD40 by cDC1 (Fig. 5D), decreased expression of CD80 and CD86 by cDC1 (Fig. 5C), and decreased expression of CD80 by cDC2 (Fig. 5B).

To determine if the changes in cDCs from Fc.Mut24-treated mice reflected transcriptional or post-transcriptional regulation, we performed bulk RNA-seq on sorted cDC1s and cDC2s from each of the four groups of mice outlined in Fig 5A. cDC1s expressed canonical genes such as *Xcr1*, *Clec9a*, *Itgae* (CD103), and *Irf8*, whereas cDC2s expressed *Irf4*, *Cd4*, and *Sirpa* (Fig. S5A). Principal-component analysis (PCA) analysis revealed that unstimulated (Ctrl) samples separated from LPS-stimulated samples across the PC1 axis for cDC1s and cDC2s indicating a strong transcriptional response to LPS in both cell types (Fig. S6A). When comparing PBS and Fc.Mut24 treatment, cDC2s clustered close together indicating that Fc.Mut24 had minimal effect on the transcriptome of cDC2s and did not alter their response to LPS. In contrast, cDC1s showed clear separation along the PC2 axis based on Fc.Mut24 treatment, with the most separation observed for LPS-stimulated samples (Fig. S6A). However, analysis of all differentially expressed genes between unstimulated and LPS-stimulated samples demonstrated that Fc.Mut24 had minimal impact on LPS-induced changes in gene expression in cDC1s, and essentially no impact in cDC2s (Fig. S6B). For cDC1s in Fc.Mut24-treated mice, a subset of 634 genes did have blunted LPS-mediated upregulation (highlighted in the red rectangle, Fig. S6B). Gene set enrichment analysis (GSEA) showed that the Hallmark gene set TNFA Signaling via NFKB was enriched in LPS-stimulated cDC1s from PBS mice compared to Fc.Mut24-treated mice (Fig. S6C). Consistent with this, cDC1s from Fc.Mut24-treated mice had slightly reduced upregulation of NF-κB-regulated genes associated with cDC maturation such as *Cd80, Il15, Il15ra, Ccr7, and Ccl5* (Fig. S6D) (*50–52*). Although *Cd80* expression was reduced in LPS-stimulated cDC1s from Fc.Mut24-treated mice, there were no changes in *Cd86* (Fig. S6E) Additionally, genes encoding MHC class II (*H2-Ab1* and *H2-Aa*) were increased in Fc.Mut24-treated mice (Fig. S6E), in contrast to what was observed for MHC class II protein expression (Fig. 5E). Together, these data show that Fc.Mut24 treatment has a minor effect on the transcriptional response of cDC1s but not cDC2s to a strong inflammatory stimulus and suggests that regulation of CD80, CD86, and MHC class II expression by T_R_ cells mainly occurs at the protein level.

### Fc.Mut24-mediated suppression of costimulatory ligand protein expression in cDCs is CTLA-4-dependent

Whereas CTLA-4 expression by T_R_ cells has been implicated in downregulating CD80 and CD86 on APCs in various settings *in vitro* (*18, 19, 53, 54*), fewer *in vivo* studies have shown this is a CTLA-4-dependent process (*20, 55*). In addition to the CTLA-4 transendocytosis model, an alternative model of trogocytosis has also been proposed, in which CD80 and CD86 plus other membrane proteins such as CD40 and MHC class II are concomitantly transferred from APCs to the T_R_ cell surface in a CTLA-4-dependent manner (*56*). We therefore wanted to understand to what extent the downregulation of CD80, CD86, and MHC class II on cDCs we observed after Fc.Mut24 treatment was dependent on CTLA-4 expression. For this, we used the same LPS stimulation model with Fc.Mut24 treatment, but before LPS administration, we blocked CTLA-4 with the 4F10 antibody (Fig. 5F). The 4F10 clone was used because it blocks CTLA-4 function but does not cause FcR-dependent depletion of T_R_ cells as seen with other anti-CTLA-4 monoclonal antibodies (*57*), which was confirmed in our studies (Fig. S7A). Anti-CTLA-4 treatment reduced total CTLA-4 expression on Foxp3^+^ T_R_ cells (Fig. S7B), but slightly increased T_R_ percentages and ICOS expression (Fig. S7A&B), likely due to increased CD28 costimulation. As expected, treatment with Fc.Mut24 followed by isotype control antibody injection impaired the LPS-mediated upregulation of CD80, CD86, and MHC class II by DCs (Fig. 5G). However, blocking CTLA-4 in Fc.Mut24-treated mice partially restored CD80 and CD86 expression by cDC1s and cDC2s, with no effect on MHC class II (Fig. 5G). Therefore, our data are consistent with the CTLA-4 transendocytosis model in which cognate CD80 and CD86 ligands, but not other membrane proteins such as MHC class II, are stripped from cDCs in a CTLA-4-dependent manner.

### Fc.Mut24-mediated suppression of MHC class II protein expression in cDC1s is IL-10-dependent

In addition to CTLA-4 expression, IL-10 production by T_R_ may also be involved in regulating cDC function in Fc.Mut24-treated mice. We found that IL-10 production was mostly restricted to CTLA-4^hi^ T_R_ cells in both PBS controls and Fc.Mut24-treated mice (Fig. 6A). Although the percentage of IL-10^+^ T_R_ cells did not change after Fc.Mut24 treatment, the absolute number of IL-10^+^ T_R_ cells significantly increased, as well as the amount of IL-10 produced on a per-cell basis (Fig. 6B). There was also a slight but significant increase in the percentage of IL-10^+^ cells among the Foxp3^-^ T_conv_ population, but this was not reflected by an increase in cell number (Fig. 6B).

**Figure 6.**
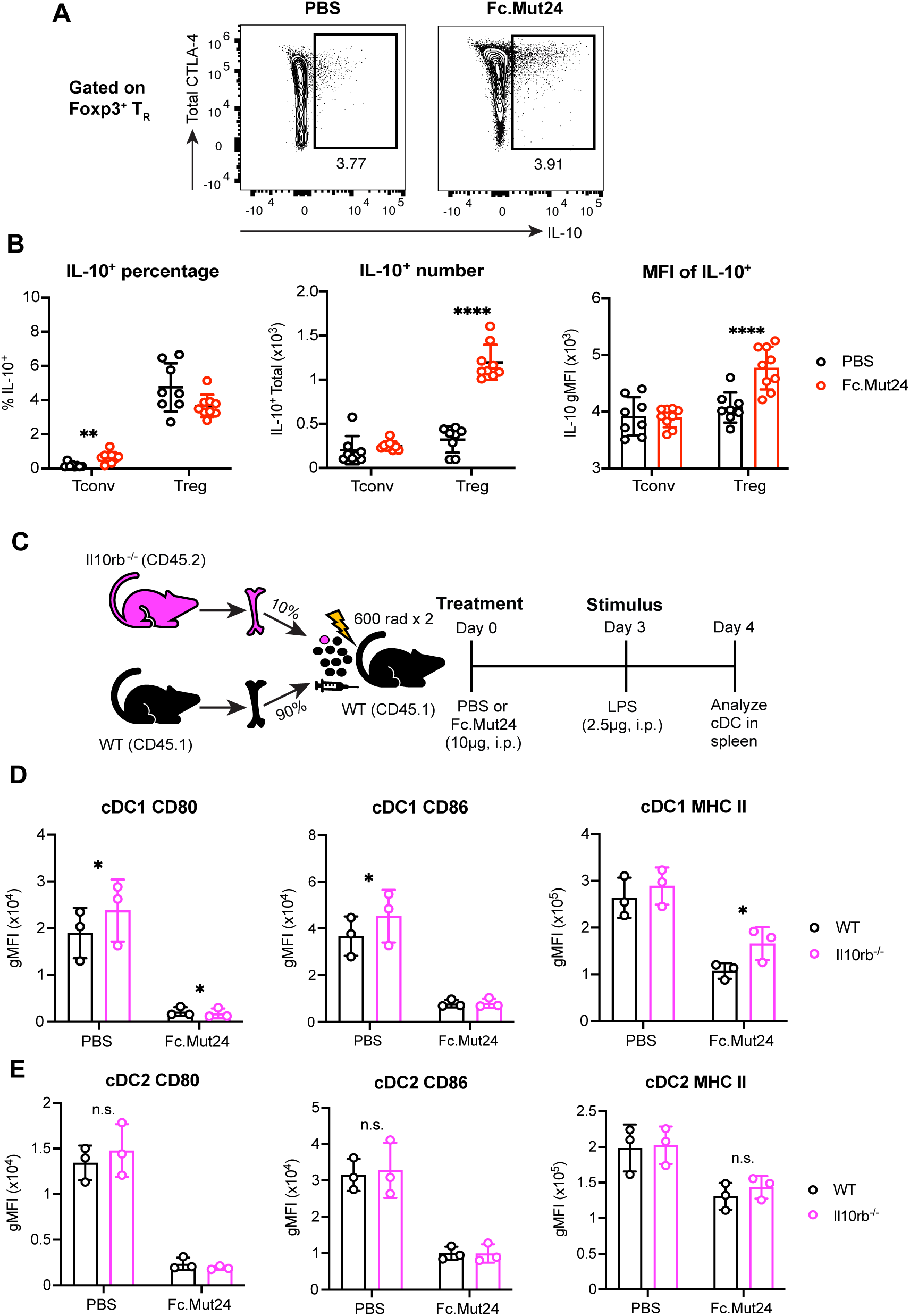
Fc.Mut24 promotes IL-10-dependent downregulation of MHC class II during dendritic cell maturation. **A&B.** Splenocytes from C57BL/6 mice treated with PBS or Fc.Mut24 were harvested 3 days later, stimulated for 2 h with PMA/Ionomycin, and analyzed by flow cytometry for IL-10 expression. Gates were set on live, TCRβ^+^, CD4^+^, Foxp3^+^ regulatory T (T_R_) or Foxp3^-^ T_conv_ cells. Representative flow cytometry plots for T_R_ are shown. **B.** Graphs show mean ± SD with individual data points (n = 8 to 9); \*\**P* ≤ 0.01, \*\*\*\**P* ≤ 0.0001, multiple unpaired *t*-tests. The mean fluorescence intensity (MFI) shown is gated on IL-10^+^ T_R_. Data are representative of 2 independent experiments. **C.** Experimental design schematic for IL-10 receptor beta knockout (*Il10rb*^-/-^) study. **D&E.** Expression of indicated DC maturation markers between wild-type and *Il10rb*^-/-^ cDC1 **(D)** and cDC2 **(E)**. Graphs show mean ± SD with individual data points (n = 3); * *P* ≤ 0.05, multiple paired *t*-tests.

To analyze the contribution of IL-10 receptor signaling to the regulation of CD80, CD86 and MHC class II in cDCs, we generated mixed bone marrow (BM) chimeric mice in which lethally irradiated CD45.1^+^ hosts were reconstituted with 90% wild-type CD45.1^+^ BM and 10% BM from CD45.2^+^ IL-10 receptor subunit beta knock-out (*Il10rb*^-/-^) mice (Fig. 6C). We used a low frequency of *Il10rb*^-/-^ cells to decrease the likelihood of recipients developing the chronic colitis or sensitivity to LPS-induced shock that can occur in global *ll10rb*^-/-^ or IL-10 knockout mice (*58, 59*). This approach allowed us to directly compare IL-10 receptor sufficient and deficient cDCs within the same animals following LPS stimulation in the absence or presence of Fc.Mut24 treatment. We found that MHC class II expression was partially restored on *Il10rb*^-/-^ compared to wild-type cDC1s from Fc.Mut24-treated mice, without any changes in CD80 or CD86 (Fig. 6D). In PBS controls, lack of IL-10 signaling did increase CD80 and CD86 levels by cDC1s, demonstrating that costimulatory ligand expression in the steady state is regulated by mechanisms other than CTLA-4-dependent transendocytosis (Fig. 6D). In contrast to cDC1s, we did not detect any significant changes in MHC class II, CD80, or CD86 expression by *Il10rb*^-/-^ versus wild-type cDC2s (Fig. 6E). IL-10 regulates MHC class II post-transcriptionally in DCs by inducing expression of the E3 ubiquitin ligase March-I that targets MHC class II molecules for degradation (*60, 61*). Consistent with a potential role for March-I in decreasing MHC class II expression in cDC1s, we found that although not statistically significant (FDR=0.065), there was a trend toward increased *March1* expression in LPS-stimulated cDC1s from mice treated with Fc.Mut24 relative to PBS controls, with minimal difference in cDC2s (Fig. S8A). Thus, during Fc.Mut24 treatment, IL-10 receptor signaling in cDC1s plays a role in downregulating MHC class II surface expression, likely through a March-1-dependent mechanism.

### Costimulatory ligand and MHC class II protein expression in cDCs is suppressed with Fc.Mut24 in autoimmune diabetes

Our prior work demonstrated that Fc.Mut24 treatment in prediabetic non-obese diabetic (NOD) mice prevents diabetes onset and reduces the severity of islet infiltration (*9*). However, the mechanisms responsible for durable disease protection have not been explored and whether distinct types of APCs are suppressed by T_R_ cells during an autoimmune response is undetermined. In NOD mice, pancreatic islet-infiltrating cDC1s and islet-resident macrophages play essential and non-redundant roles in the development of autoimmune diabetes (*62, 63*). Migratory cDC1s (XCR1^+^CD103^+^) prime islet-reactive CD8^+^ and CD4^+^ T cells in the pancreatic lymph node (pancLN), an early event in the inflammatory cascade, and present autoantigens to T cells in the islets during disease progression (*62*), whereas resident macrophages (F4/80^+^) promote the early entrance of CD4^+^ T cells and cDCs into the islets (*63*).

As previously shown, treatment of prediabetic NOD mice with Fc.Mut24 significantly increased the percentage and number of Foxp3^+^ T_R_ cells in the spleen, pancLN, and pancreas, with almost all cells being Ki-67^+^ (Fig. S9A). Similar to what was observed for splenic T_R_ cells in B6 mice, Fc.Mut24 treatment in NOD mice led to the upregulation of CD25, Foxp3, CTLA-4, and ICOS in splenic, pancLN, and pancreatic T_R_ cells and the downregulation of PD-1 (Fig. S9B). To establish how T_R_ expansion and activation affect disease-relevant APCs, we analyzed XCR1^+^CD103^+^ migratory cDC1s and Sirpα^+^ cDC2s in the pancLN and pancreas, as well as pancreatic F4/80^+^ macrophages in PBS controls or Fc.Mut24-treated mice (Fig. S10A). There was an expansion of XCR1^+^ cDC1 number in the spleen, pancLN, and pancreas after Fc.Mut24 treatment and a slight increase in splenic Sirpα^+^ cDC2 number, but no changes in pancreatic F4/80^+^ macrophages (Fig. S10B). In agreement with reports showing a pro-inflammatory phenotype (*62–64*), pancreatic F4/80^+^ macrophages had high expression CD80, CD86, CD40, and MHC class II (Fig. 7A), and treatment with Fc.Mut24 did not alter the expression of any of these markers by macrophages (Fig. 7A). In contrast, CD80 expression was significantly reduced in migratory cDC1s from the pancLN and pancreas after treatment (Fig. 7A). CD86 expression was also lower in pancreatic cDC1s from Fc.Mut24-treated mice, in addition to lower expression of CD80 and MHC class II in pancreatic cDC2s (Fig. 7A). Across all cell types and tissues examined, CD40 was not downregulated by Fc.Mut24 treatment (Fig. 7A). Together, these data demonstrate that T_R_ cells in lymphoid tissues and at the site of tissue inflammation are activated with Fc.Mut24, leading to complex tissue- and cell-type-dependent effects on cDCs but not macrophages.

**Figure 7.**
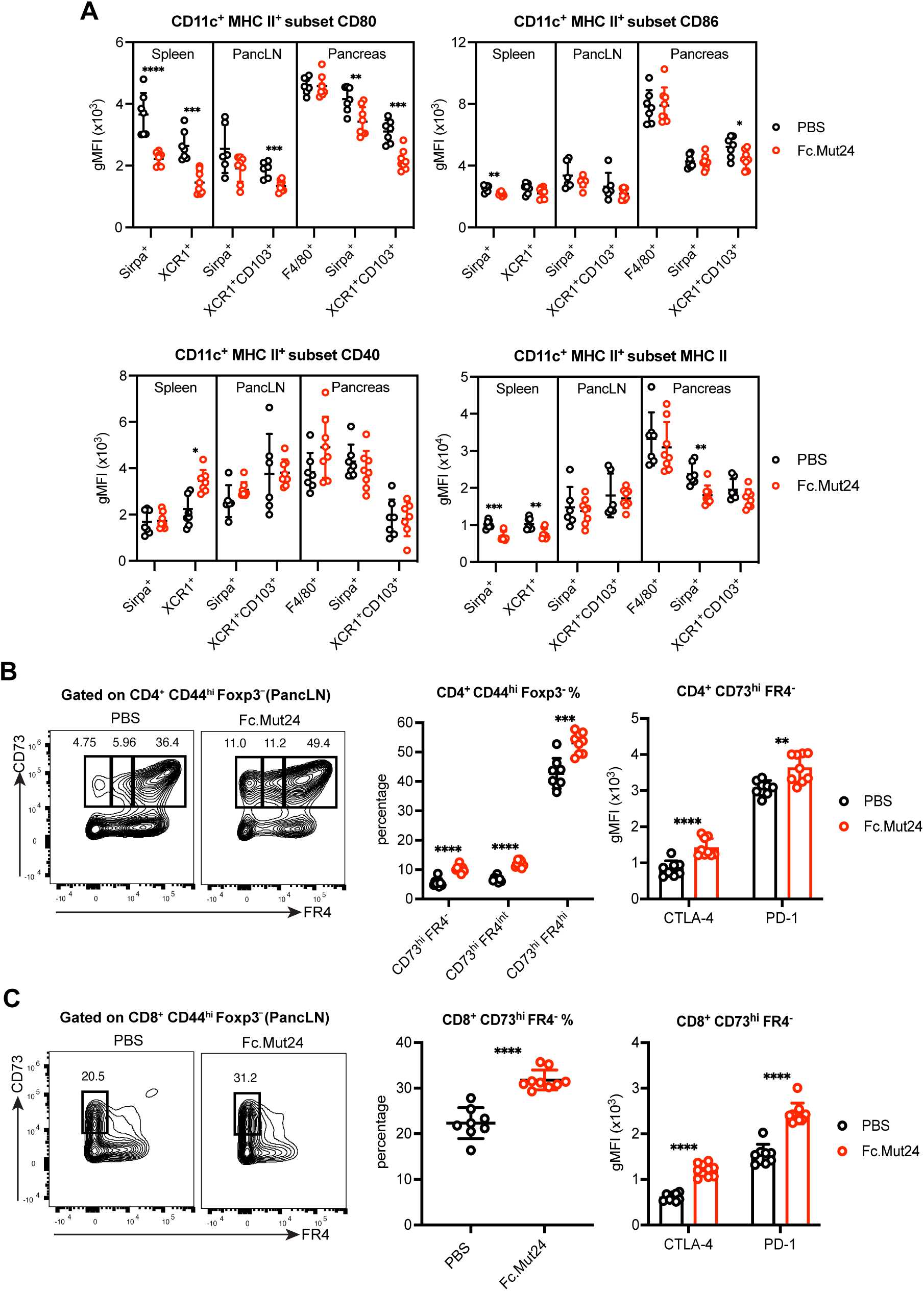
Fc.Mut24 limits dendritic cell maturation and induces anergic T cells during the progression of autoimmune diabetes. **A.** Expression of indicated maturation markers on different subsets of CD11c^+^ MHC II^+^ antigen-presenting cells isolated from the spleen, pancreatic lymph node (PancLN), and pancreas of 8-10-week-old female non-obese diabetic (NOD) mice treated with PBS or Fc.Mut24 4 days previously. See Fig. S10A for the gating strategy. **B.** Percentage and number of CD73^hi^ FR4^-^, CD73^hi^ FR4^intermediate^, and CD73^hi^ FR4^hi^ conventional T (T_conv_) cells isolated from the PancLN and expression of indicated markers on CD73^hi^ FR4^-^ cells. Gates were set on live, CD45^+^, CD4^+^, CD44^hi^, Foxp3^-^ cells. **C.** Percentage and number of CD73^hi^ FR4^-^ T_conv_ cells isolated from the PancLN and expression of indicated markers. Gates were set on live, CD45^+^, CD8^+^, CD44^hi^, Foxp3^-^ cells. **A-C.** All graphs show mean ± SD with individual data points (n = 6 to 9); \**P* ≤ 0.05, \*\**P* ≤ 0.01, \*\*\**P* ≤ 0.001, \*\*\*\**P* ≤ 0.0001, two-tailed unpaired t-tests, or multiple unpaired *t*-tests. Data are representative of 2-3 independent experiments.

### Induction of anergic T cells with Fc.Mut24 in autoimmune diabetes

Since the upregulation of CTLA-4 by Foxp3^+^ T_R_ cells in the pancLN of Fc.Mut24-treated NOD mice correlated with decreased CD80 expression by migratory cDC1s we wanted to determine the impact on anergy in T_conv_ cells at this site. We first analyzed CD4^+^ CD44^hi^ Foxp3^-^ cells for co-expression of CD73 and FR4, which defines an anergic population that has the ability to differentiate into Foxp3^+^ T_R_ cells (*65*). Differentiation of anergic CD4^+^ T cells into suppressive Foxp3^-^ Tr1 cells by tolerogenic DCs has also been reported and is associated with the upregulation of CD73, CTLA-4, and PD-1 (*66, 67*). We found that the percentage and number of anergic CD73^hi^ FR4^hi^ cells as well as CD73^hi^ FR4^intermediate^ and CD73^hi^ FR4^-^ cells in the pancLN significantly increased after Fc.Mut24 treatment (Fig. 7B). The CD73^hi^ FR4^-^ population also had increased expression of CTLA-4 and PD-1 in Fc.Mut24-treated mice (Fig. 7B), consistent with what was previously reported for induced Foxp3^-^ Tr1 cells. When analyzing CD8^+^ CD44^hi^ Foxp3^-^ cells, we observed a similar expansion of CD73^hi^ FR4^-^ cells after Fc.Mut24 treatment in addition to the upregulation of CTLA-4 and PD-1 by this population (Fig. 7C). These findings suggest that by reducing costimulatory ligand expression on cDC1s in the pancLN, Fc.Mut24-expanded Foxp3^+^ T_R_ cells promote anergy in autoreactive T_conv_, with the potential for generating suppressive CD4^+^ and CD8^+^ T cells lacking Foxp3 expression that may also contribute to immune regulation as an example of infectious tolerance (*68*).

## Discussion

Despite the current testing of CD25-biased Fc.IL-2 muteins with enhanced Foxp3^+^ T_R_ cell selectivity in clinical trials (*10*), the molecular mechanisms by which these therapeutics induce immune tolerance and prevent disease progression in models of autoimmune disease have not been well characterized. In addition, although the role of IL-2 in maintaining the homeostasis of Foxp3^+^ T_R_ cells is well documented (*21, 69*), and there is an indispensable role of the IL-2 receptor (IL-2R) in T_R_ suppressor activity (*37*), there is still a lack of complete understanding of how IL-2R signaling influences T_R_ differentiation and function.

We used a murine Fc.IL-2 mutein (Fc.Mut24) that we previously developed for preclinical studies as a tool to understand how treatment influences Foxp3^+^ T_R_ suppressive function *in vivo*. We found that Fc.Mut24 treatment induced a large expansion of central effector T_R_ (ceT_R_) cells with high expression of the activation marker CD44 and the lymphoid trafficking molecule CD62L, increased expression of immunoregulatory molecules such as CTLA-4 and IL-10, and decreased expression of molecules that antagonize T_R_ function, like PD-1. Similar to our findings, low-dose IL-2 therapy promotes the upregulation of CD44 and downregulation of PD-1 on antigen-specific CD8^+^ T cells during chronic viral infection and is associated with enhanced effector function (*70*). Although the origin of the ceT_R_ population after Fc.Mut24 treatment will require further investigation, these cells may derive from central T_R_ (cT_R_) that upregulate CD44 or effector T_R_ (eT_R_) that upregulate CD62L. Neither *Cd44* nor *Sell* (the gene encoding CD62L) appear to be direct targets of Stat5 in T_R_, yet both are modulated by TCR signaling and their altered expression in Fc.Mut24-treated mice could reflect prolonged IL-2R and TCR synergy due to the enhanced half-life and signaling of Fc.Mut24. Consistent with this hypothesis, we found that Fc.Mut24 treatment induced a distinct transcriptional profile indicative of both IL-2R and TCR signaling and a recent report showed that T_R_ cell expansion with Fc.IL-2 muteins was blunted by the blockade of MHC class II (*71*). Whether unique phenotypes are observed in human Foxp3^+^ T_R_ after Fc.IL-2 mutein treatment will be interesting to examine, given the heterogeneity in human T_R_ populations that we and others have described (*72, 73*).

Our data demonstrate that a major mechanism by which Fc.Mut24 treatment alters T_R_ function is through increasing CTLA-4 expression and cycling from intracellular compartments to the cell surface in Foxp3^+^ T_R_ but not Foxp3^-^ conventional T (T_conv_) cells. Previous work has pointed to TCR signaling as the principal regulator of CTLA-4 function, as T cell activation via anti-CD3 increases CTLA-4 cycling in both T_R_ and T_conv_ cells, albeit with different kinetics between cell types (*20, 43*). However, these experiments were performed using total CD4^+^ T cells rather than purified populations, and it cannot be ruled out that IL-2 produced by T_conv_ upon activation synergized with TCR signaling to enhance CTLA-4 cycling in T_R_ cells. A connection between IL-2-induced immune regulation and the CTLA-4 pathway was previously suggested since IL-2 induces the surface and intracellular expression of CTLA-4 in murine and human T cells *in vitro* (*74, 75*), and acute IL-2 blockade in mice decreases CTLA-4 expression by T_R_ cells (*76*). Transgenic expression of CTLA-4 also rescues the lymphoproliferative disease that occurs in IL-2-deficient mice (*77*), suggesting that impaired CTLA-4 function in the absence of IL-2 contributes to immune pathology. In support of our CTLA-4 cycling results, Fc.Mut24 treatment significantly increased CTLA-4-dependent transendocytosis of CD86 *ex vivo* and CD80 *in vivo* by T_R_ but not T_conv_ cells. We were particularly interested to find that cT_R_ cells, which usually have a naïve phenotype and relatively low CTLA-4 expression, gained the ability to capture CD86 in Fc.Mut24-treated mice. As cT_R_ are generally found at a high density within T cell zones of secondary lymphoid tissues where T_conv_-dendritic cell (DC) interactions and T cell priming occur, this increases their potential to contribute to immune regulation after Fc.Mut24 treatment.

To determine if CTLA-4 upregulation by T_R_ after Fc.Mut24 treatment resulted in the suppression of conventional DC (cDC) function, we established a model of LPS-mediated maturation. In these experiments and others in non-obese diabetic (NOD) mice, Fc.Mut24 increased the abundance of cDC1s in multiple tissues. While we did not further explore this, IL-2 has been shown to induce cDC expansion by increasing T cell production of cytokines such as Fms-like tyrosine kinase 3 ligand (Flt3l) (*78*). Importantly, Fc.Mut24 significantly enhanced CTLA-4-dependent transendocytosis of CD80 and CD86 during splenic cDC1 and cDC2 maturation, in addition to downregulation of surface MHC class II, which was not CTLA-4-dependent but due to IL-10 signaling in cDC1s. Despite the strong effect of Fc.Mut24 treatment on CD80 and CD86 protein expression, treatment slightly blunted the overall transcriptional response of cDC1s but not cDC2s to LPS stimulation. While the reasons for the differential transcriptional regulation of splenic cDC1 versus cDC2 subsets are not apparent, at steady state cDC1s are found within splenic T cell zones, whereas cDC2s are located at the marginal zone bridging channel between the red pulp and white pulp (*79*), possibly allowing for more sustained interactions between T_R_ cells and cDC1s. In human monocytes, IL-10 prevents IκB degradation as well as nuclear translocation of NF-κB and binding to target genes (*80*). Therefore, IL-10-mediated downregulation of the NF-κB pathway may reduce CD80 transcription in cDC1s and somewhat blunt their transcriptional response to LPS. We found no evidence of CD40 downregulation during cDC maturation, similar to other studies on T_R_ cells activated via TCR stimulation, which downregulate CD80 and CD86, but not CD40, on splenic DCs (*54*). These results support a model in which increased CTLA-4-dependent transendocytosis and IL-10 production are important mechanisms used by T_R_ to modulate cDC function following Fc.Mut24 treatment, and establish that in the presence of a strong inflammatory stimulus, T_R_ cells do not completely inhibit cDC maturation.

In cancer models, T_R_ cells suppress cDC1s in both the tumor-draining lymph node and the tumor (*81, 82*), but it is unclear whether similar findings apply to target organs in autoimmune disease. Using 8-10 week-old female NOD mice, an age at which there is already established autoimmunity and infiltration of the pancreatic islets but no overt diabetes (*83*), we found that cDC2s and migratory cDC1s from the pancreas of Fc.Mut24-treated mice had reduced CD80 expression and there was reduced CD86 expression by cDC1s. These migratory cDC1s, which are required for the initiation of disease development in NOD mice (*62*), also had lower CD80 expression in the pancreatic lymph node. Consistent with this, the frequency of anergic CD73^hi^ FR4^hi^ CD4^+^ T cells in the pancreatic lymph node and CD73^hi^ FR4^-^ T cells that were either CD4^+^ or CD8^+^ significantly increased after Fc.Mut24 treatment. In CD8^+^ T cells, a lack of CD28 costimulation during activation leads to CD73 upregulation and the generation of regulatory cells that suppress effector T cell responses through CD73-mediated adenosine production (*84*). Similarly, CD4^+^ T cells activated without CD28 costimulation have elevated levels of the *Nt5e* gene encoding CD73 (*85*), suggesting that CD73 upregulation is a common mechanism of T cell tolerance in the absence of sufficient costimulation.

High levels of IL-2R signaling promote the terminal differentiation of short-lived but highly suppressive T_R_ cells (*86*). This is analogous to the ability of strong IL-2R signaling to induce terminally differentiated short-lived effector CD8^+^ T cells (*2*), and highlights a conserved role for IL-2R signaling driving both T_R_ and T_conv_ effector function. While the mechanisms by which strong IL-2R signaling potentiates T_R_ suppressive function are not completely understood, Stat5 activation appears to increase T_R_ adherence to DCs *in vitro*, reducing their expression of CD80 and CD86 (*37*). Exposure of T_R_ cells to IL-2 also increases T_R_-DC interactions *in vivo,* leading to suppression of DC function and T cell priming in a contact-dependent but MHC class II-independent manner (*87*). We postulate that extensive IL-2R signaling in T_R_ cells increases CTLA-4 expression and T_R_-DC adhesion, allowing T_R_ to strip DCs of CD80 and CD86 in an MHC class II-independent manner, which is consistent with the established models of T_R_-mediated bystander suppression.

In conclusion, evidence from this study and others demonstrates that CD25-biased Fc.IL-2 muteins have unique properties such as sustained IL-2 signaling that promote the expansion and differentiation of highly activated Foxp3^+^ T_R_ cells. We show that these Fc.IL-2 mutein-expanded T_R_ cells have a distinct transcriptional and functional profile and can regulate DCs using multiple molecular mechanisms resulting in reduced costimulatory ligand availability that is accompanied by induction of T cell anergy and expression of the immunoinhibitory protein CD73 by T_conv_ cells. The relationship between sustained IL-2R signaling, T_R_ effector function, and anergy induction is highly desirable under conditions of persistent autoreactive T cell activation for establishing lasting immune tolerance. Indeed, the ability of Fc.IL-2 muteins to enhance T_R_-mediated suppression of DCs during an autoimmune response distinguishes this class of therapeutics from other biologics that block pathogenic effector T cell pathways, further warranting their use in the prevention and treatment of chronic inflammatory diseases.

## Supporting information

Key Resources Table

## Abbreviations

IL-2: Interleukin-2
mutein: mutant protein
IL-10: Interleukin-10
TCR: T cell receptor
CTLA-4: Cytotoxic T-lymphocyte Antigen 4
NOD: non-obese diabetic
APC: antigen-presenting cell
cDC: conventional dendritic cell
T_conv_: conventional T cell
T_R_: regulatory T cell
cT_R_: central T_R_
ceT_R_: central effector T_R_
eT_R_: effector T_R_;
LPS: Lipopolysaccharide.

## Acknowledgments

We thank Shivani Srivastava (Fred Hutchinson Cancer Center) for the gift of the Plat-E packaging cells, Francesco Marangoni (UC Irvine) for help with the signal amplification protocol used for the staining of surface and cycling CTLA-4, the Cell and Tissue Analysis Group at the Benaroya Research Institute (BRI) for flow cytometry and sorting expertise with assistance from Adam Wojno and Aiden Pierce, and the Genomics Core at BRI for bulk and single-cell RNA sequencing with assistance from Vivian Gersuk.

## Author contributions

B.L.J. designed and performed the experiments, analyzed and interpreted the data, prepared the figures, and co-wrote the manuscript. L.S.K.W. and D.M.S. generated the CD80-mCherry mice. C.J.W. and L.S.K.W. designed and performed experiments on *in vivo* CTLA-4 transendocytosis. C.J.W., L.S.K.W, and M.A.G. edited the manuscript. M.L. and H.D. analyzed the data and assisted with figure generation. D.J.C. and M.A.G. conceptualized this study, and D.J.C. supervised the study and co-wrote the manuscript.

## Methods

### Mice

C57BL/6 (B6), C57BL/6.CD45.1^+^ (JAXBoy, B6.CD45.1^+^), C57BL/6.CRFB4^-/-^ (B6.Il10rb^-/-^) and NOD/ShiLtJ (NOD) mice were purchased from The Jackson Laboratory. C57BL/6.Foxp3-mRFP (B6.Foxp3-mRFP) mice were originally purchased from the Jackson Laboratory and bred in-house. The B6.Foxp3-mRFP mice have a monomeric red fluorescent protein (mRFP) knocked in downstream of Foxp3. Mixed bone marrow (BM) chimeric mice were made by transferring 0.5x10^6^ B6.Il10rb^-/-^ (CD45.2^+^) BM cells and 4.5x10^6^ B6.CD45.1^+^ BM cells intravenously into lethally irradiated B6.CD45.1^+^ hosts. Mixed BM chimeric mice were allowed to reconstitute for at least 9 weeks. NOD mice were confirmed to be prediabetic (normal blood glucose) at the time of analysis. The B6, B6.CD45.1, B6.Il10rb^-/-^, B6.Foxp3-mRFP, and NOD mice were maintained at the Benaroya Research Institute (BRI, Seattle, WA), and experiments were performed in accordance with the guidelines of the Institutional Animal Care and Use Committee of BRI. Mice used in experiments at BRI were between 8 and 12 weeks of age.

CD80-mCherry.Rag2^-/-^ mice were generated and bred in-house at University College London (UCL, London, UK). CD80-mCherry was expressed under the control of the endogenous CD80 promoter, and mice were generated on a BALB/c background (Wang et al., manuscript in preparation). Wild-type BALB/c mice were originally purchased from the Jackson Laboratory and bred in-house at UCL. For experiments at UCL, mice were housed in individually ventilated cages (IVC) with environmental enrichment (e.g., cardboard tunnels, paper houses, chewing blocks, and aspen wood wool nesting material) in a temperature- and humidity-controlled facility with a 14-h light/10-h dark cycle and ad libitum feeding. Experimental animals were located in the middle rows of the IVC rack to minimize the impact of differences in light exposure. All injections were carried out in the absence of anesthesia and analgesia, typically in the morning, and mice were returned to the home cage immediately following the procedure. The first injection for *in vivo* CTLA-4 transendocytosis experiments was typically between 3 pm and 6 pm. The welfare of experimental animals was monitored regularly (typically immediately post-procedure, then at least every 2–3 days) and no procedure-related adverse events were noted. For the majority of experiments, animals were randomly assigned into treatment groups after being matched for age and no blinding was used. No data points were excluded. The number of replicates is provided in the figure legend. Mice used in experiments at UCL were between 12 and 14 weeks of age.

### Fc.IL-2 mutein

The development of the murine Fc.IL-2 mutein (Fc.Mut24) and Fc.WT IL-2 was previously described (*9*). These molecules were produced and purified by Olympic Protein Technologies (Seattle, WA). Purified Fc.Mut24 and Fc.WT contained less than 15 endotoxin units (EU)/ml. For all experiments, 10 µg of Fc.WT or Fc.Mut24 was administered via intraperitoneal injection.

### Single-cell RNA-seq of Foxp3^+^ T_R_ cells

CD4^+^ T cells from the spleen of Foxp3-mRFP mice were enriched using a CD4 negative selection kit (Miltenyi Biotec) according to the manufacturer’s protocol. Enriched CD4^+^ T cells were stained with viability dye and labeled with anti-mouse CD16/32 (Biolegend). Samples were stained with TotalSeq-C anti-mouse hashtag antibodies (Biolegend), and 1 x 10^5^ CD4^+^ Foxp3^+^ T_R_ per sample were sorted (FACS Aria II, Becton Dickinson) into RPMI-10. Samples were pooled in groups of 3 for a total of four pooled samples (n = 12). Lineage antibodies to exclude populations during sorting included those targeting CD8 (cytotoxic T cells), CD19 (B cells), GR-1 (neutrophils), and NK1.1 (NK cells). A single cell suspension from each pooled sample was loaded in a single channel of the 10x Chromium Controller (10X Genomics). Sequencing libraries were generated using the NextGEM Single Cell 5’ Kit v2 kit. Gene expression, mouse TCR, and feature barcoding libraries were pooled and treated with Illumina Free Adapter Blocking Reagent (Illumina). Sequencing of pooled libraries was carried out on a NextSeq 2000 sequencer (Illumina), using two NextSeq P3 flowcells (Illumina) with the aim of capturing 1.5 x 10^4^ cells per pooled sample and a target sequencing depth of 30,000/reads per cell.

### Demultiplexing and alignment of single-cell RNA-seq data

Cell Ranger (version 6.1.1, 10x Genomics) mkfastq was used to demultiplex and produce raw fastq files for downstream analyses. Cell Ranger multi was used to align per-pool reads against the reference mouse transcriptome (mm10), in addition to barcoded hash tags, Cell Ranger vdj was called to assign TCR reads to cells. All three assays were aggregated across the four pools using Cell-Ranger aggr to produce raw count matrices for downstream analyses. Cells containing fewer than 500 genes were removed from this matrix, as well as any cell exceeding 4,500 features, with 12.5% and 5% of genes assigned to mitochondria and hemeglobin genes, respectively. Expression data of all assays were normalized (RNA: log-norm) and scaled to produce final matrices for downstream analyses.

### Analysis of single-cell RNA-seq data

Principle component analysis (PCA) of RNA expression was performed in Seurat 2 (*88*). The top 30 PCs from RNA were reduced into UMAP space for visualization, followed by Louvain clustering in Seurat for cluster assignment. Top genes for all clusters and cell identities reported in this study were calculated by Wilcoxon rank sum tests within Seurat to identify all genes with p < .05. The average expression of these genes across clusters was determined by averageExpression in Seurat, followed by scaling to convert to z-scores. Heatmaps of these z-scores were produced via the package ComplexHeatmap. Gene module scores for modules of interest were assigned via addModuleScore within Seurat, and cells were manually gated based on module scores into high- and low-scoring cells in order to produce Violin Plots. All plots were produced using ggplot2. All statistical analyses were performed in R version 4.2.1.

### Tissue preparation for flow cytometry and cDC sorting

To isolate T cells from B6 mice, spleen samples were mashed through 70-um strainers into RPMI 1640 plus 10% FBS (RPMI-10). To isolate CD11c^+^ cells from B6 mice, minced whole spleens were digested in basal RPMI 1640 supplemented with 25 ug/ml Liberase TM (Roche) and 25 ug/ml DNase I (Sigma-Aldrich) for 20 min under agitation at 37°C. Erythrocytes were lysed in ACK lysis buffer, and the remaining cells were washed with RPMI-10. CD11c^+^ cells were enriched using CD11c MicroBeads (Miltenyi Biotec) according to the manufacturer’s protocol. To isolate T cells and CD11c^+^ cells from non-obese diabetic (NOD) mice, minced whole spleens and pancreatic lymph nodes (PancLN) were digested in basal RPMI 1640 supplemented with 25 ug/ml Liberase TM (Roche) and 25 ug/ml DNase I for 20 min under agitation at 37°C. Erythrocytes were lysed in ACK lysis buffer for spleen samples. Minced pancreas samples from NOD mice were digested in RPMI-10 supplemented with 5 mg/ml Collagenase Type V (Sigma) and 10 ug/ml DNase I followed by incubation in enzyme-free cell dissociation buffer (Sigma) for 5-10 min at room temperature. CD11c^+^ cells were enriched from the spleen and pancreas, but not pancLN samples of NOD mice, using CD11c MicroBeads (Miltenyi Biotec) according to the manufacturer’s protocol.

### Flow Cytometry

For flow cytometric analysis, cells were stained with anti-mouse CD16/32 (Biolegend) for 10 min at 4°C to block nonspecific binding to Fc receptors, followed by cell surface staining in FACS buffer (PBS-2% BCS) for 20-30 min at 4°C. For staining CD11c^+^ samples, lineage antibodies included those targeting CD5 (T cells), CD19 (B cells), GR-1 (neutrophils), and NK1.1 (NK cells). For intracellular antigens, cells were fixed and permeabilized using the Foxp3/Transcription Factor Staining Buffer Set (eBioscience), followed by staining in permeabilization buffer for 20 min at room temperature. Before fixation, dead cells were labeled using Fixable Viability Dye eFluor780 (ThermoFisher). For measuring IL-10 expression, intracellular staining was performed for 2 h at 4°C after stimulation for 2 h with PMA/Ionomycin Cell Stimulation Cocktail containing protein transport inhibitors (ThermoFisher). Data were acquired on an Aurora (Cytek Biosciences) or Symphony (BD Biosciences) flow cytometer and analyzed with FlowJo (FlowJo LLC). Counting beads were added to samples to quantify cell numbers.

### *Ex vivo* CTLA-4 cycling

Splenocytes from B6 mice were labeled with Allophycocyanin (APC)-conjugated anti-CTLA-4 antibody for 30 min at 4°C to detect surface CTLA-4, for 2 h at 37°C to detect cycling CTLA-4, or after fixation and permeabilization to detect total CTLA-4. The surface CTLA-4 and cycling CTLA-4 samples were washed with FACS buffer, and secondary staining was performed with biotinylated anti-APC antibody for 20 min at 4°C, followed by tertiary staining with Streptavidin-APC for 20 min at 4°C. The signal amplification approach was used to enhance the detection of CTLA-4 antigen, which is poorly expressed on the cell surface due to its highly endocytic properties.

### Bulk RNA-seq of cDCs

300 cells per population were sorted (FACS Aria II, Becton Dickinson) into lysis buffer, and cDNA was prepared using the SMART-Seq Ultra Low Input RNA Kit for Sequencing (Takara Bio). RNA-seq libraries were constructed using the NexteraXT DNA Library Preparation Kit (Illumina) with half the recommended volumes and reagents. Paired-end sequencing of pooled libraries was run on a NextSeq 2000 (Illumina) with 59-base reads and a target depth of 5 million reads per sample. After the run, base-calling and demultiplexing were performed automatically on BaseSpace (Illumina) to generate FASTQ files. The FASTQs were aligned to the University of California Santa Cruz (UCSC) Human Genome assembly version 19, using STAR v.2.4.2a, and gene counts were generated using htseq-count. QC and metrics analysis was performed using the Picard family of tools (v1.134). To detect differentially expressed genes between cell populations, the RNA-seq analysis functionality of the linear models for microarray data (Limma) R package was used, and the ROAST method within the Limma R package was used to perform gene set enrichment analysis (*89, 90*). Expression counts were normalized using the TMM algorithm (*91*). A false discovery rate adjustment was applied to correct for multiple testing.

### *In vivo* Ab treatment

For blocking CTLA-4 *in vivo*, 750 μg of anti-CTLA-4 antibody clone UC10-4F10-11 (Bio X Cell) or hamster IgG isotype control antibody (clone PIP, Bio X Cell) was administered via intravenous injection on Day 2 after PBS or Fc.Mut24 treatment.

### *Ex vivo* CTLA-4 transendocytosis

Human CD86 C-terminally tagged with enhanced GFP (pCD86-EGFP, Addgene) was sub-cloned into a modified version of the MSCV2.2 retroviral plasmid in which the IRES-GFP cassette was removed. This plasmid was transfected into Plat-E packaging cells to produce retrovirus that was used to transduce NIH/3T3 fibroblasts. The resultant NIH/3T3 transfectants were sorted (FACS Aria Fusion, Becton Dickinson) for uniform GFP expression. CD4^+^ T cells from the spleen of B6 mice were enriched using CD4 MicroBeads (Miltenyi Biotec) according to the manufacturer’s protocol. Enriched CD4^+^ T cells were co-cultured with CD86-GFP expressing NIH/3T3 cells at a 1:1 ratio for 2 h at 37°C. Where indicated, 100 µg/ml of anti-CTLA-4 antibody (clone UC10-4F10-11, Bio X Cell) or 25 nM Bafilomycin A1 (Sigma-Aldrich) was added to the culture. After incubation, NIH/3T3 cells remained adherent while CD4^+^ T cells were removed, stained, and analyzed by flow cytometry.

### *In vivo* CTLA-4 transendocytosis

CD80-mCherry.Rag2^-/-^ mice (12-14 weeks old, male and female) were injected intravenously on Day 0 with 8x10^6^ CD4^+^ T cells purified from wild-type BALB/c lymph nodes. 500 µg of anti-CTLA-4 antibody (clone UC10-4F10-11, BioXCell) or hamster IgG isotype control antibody (clone PIP, Bio X Cell) was administered via intraperitoneal injection on Day 0, Day 3, and Day 6. PBS or Fc.Mut24 was also administered via intraperitoneal injection on Day 3. Spleen cells were analyzed by flow cytometry on Day 7.

### Statistical analysis

Statistical analysis was performed using GraphPad Prism version 9. When comparing two groups, *P* values were calculated by two-tailed Student’s *t*-tests, while multiple *t*-tests were used for comparing more than two groups. For multiple *t*-tests, the discovery was determined using the two-stage step-up method of Benjamini, Krieger, and Yekutieli, with Q = 1%. No formal sample size calculations were performed, and data distribution was assumed normal but not formally tested. The sample size, *P* values, number of replicates, and statistical tests used in each experiment are listed in the Figure legends. In animal experiments, the experimental unit is an individual mouse.

## References

1. B. H. Nelson, IL-2, regulatory T cells, and tolerance. J Immunol 172, 3983–3988 (2004).

2. V. Kalia, S. Sarkar, Regulation of Effector and Memory CD8 T Cell Differentiation by IL-2-A Balancing Act. Front Immunol 9, 2987 (2018).

3. M. Pepper, A. J. Pagán, B. Z. Igyártó, J. J. Taylor, M. K. Jenkins, Opposing signals from the Bcl6 transcription factor and the interleukin-2 receptor generate T helper 1 central and effector memory cells. Immunity 35, 583–595 (2011).

4. D. Klatzmann, A. K. Abbas, The promise of low-dose interleukin-2 therapy for autoimmune and inflammatory diseases. Nat Rev Immunol 15, 283–294 (2015).

5. H. T. Kim et al., Organ-specific response after low-dose interleukin-2 therapy for steroid-refractory chronic graft-versus-host disease. Blood Adv 6, 4392–4402 (2022).

6. J. He et al., Efficacy and safety of low-dose IL-2 in the treatment of systemic lupus erythematosus: a randomised, double-blind, placebo-controlled trial. Ann Rheum Dis 79, 141–149 (2020).

7. J. Y. Humrich et al., Low-dose interleukin-2 therapy in active systemic lupus erythematosus (LUPIL-2): a multicentre, double-blind, randomised and placebo-controlled phase II trial. Ann Rheum Dis 81, 1685–1694 (2022).

8. L. B. Peterson et al., A long-lived IL-2 mutein that selectively activates and expands regulatory T cells as a therapy for autoimmune disease. J Autoimmun 95, 1–14 (2018).

9. L. Khoryati, et al., An IL-2 mutein engineered to promote expansion of regulatory T cells arrests ongoing autoimmunity in mice. Sci Immunol 5, (2020).

10. M. E. Raeber, D. Sahin, U. Karakus, O. Boyman, A systematic review of interleukin-2-based immunotherapies in clinical trials for cancer and autoimmune diseases. EBioMedicine 90, 104539 (2023).

11. V. Lykhopiy, V. Malviya, S. Humblet-Baron, S. M. Schlenner, “IL-2 immunotherapy for targeting regulatory T cells in autoimmunity”. Genes Immun 24, 248–262 (2023).

12. B. L. Jamison et al., Nanoparticles Containing an Insulin-ChgA Hybrid Peptide Protect from Transfer of Autoimmune Diabetes by Shifting the Balance between Effector T Cells and Regulatory T Cells. J Immunol 203, 48–57 (2019).

13. R. H. Schwartz, T cell anergy. Annu Rev Immunol 21, 305–334 (2003).

14. T. L. Vanasek, S. L. Nandiwada, M. K. Jenkins, D. L. Mueller, CD25+Foxp3+ regulatory T cells facilitate CD4+ T cell clonal anergy induction during the recovery from lymphopenia. J Immunol 176, 5880–5889 (2006).

15. R. J. Martinez et al., Arthritogenic self-reactive CD4+ T cells acquire an FR4hiCD73hi anergic state in the presence of Foxp3+ regulatory T cells. J Immunol 188, 170–181 (2012).

16. C. Haase, T. N. Jørgensen, B. K. Michelsen, Both exogenous and endogenous interleukin-10 affects the maturation of bone-marrow-derived dendritic cells in vitro and strongly influences T-cell priming in vivo. Immunology 107, 489–499 (2002).

17. S. Avdic et al., Human cytomegalovirus interleukin-10 polarizes monocytes toward a deactivated M2c phenotype to repress host immune responses. J Virol 87, 10273–10282 (2013).

18. K. Wing et al., CTLA-4 control over Foxp3+ regulatory T cell function. Science 322, 271–275 (2008).

19. O. S. Qureshi et al., Trans-endocytosis of CD80 and CD86: a molecular basis for the cell-extrinsic function of CTLA-4. Science 332, 600–603 (2011).

20. V. Ovcinnikovs, et al., CTLA-4-mediated transendocytosis of costimulatory molecules primarily targets migratory dendritic cells. Sci Immunol 4, (2019).

21. K. S. Smigiel et al., CCR7 provides localized access to IL-2 and defines homeostatically distinct regulatory T cell subsets. J Exp Med 211, 121–136 (2014).

22. K. H. Toomer et al., Developmental Progression and Interrelationship of Central and Effector Regulatory T Cell Subsets. J Immunol 196, 3665–3676 (2016).

23. G. M. Wiedemann et al., Divergent Role for STAT5 in the Adaptive Responses of Natural Killer Cells. Cell Rep 33, 108498 (2020).

24. H. Nakajima et al., An indirect effect of Stat5a in IL-2-induced proliferation: a critical role for Stat5a in IL-2-mediated IL-2 receptor alpha chain induction. Immunity 7, 691–701 (1997).

25. E. P. Consortium, The ENCODE (ENCyclopedia Of DNA Elements) Project. Science 306, 636–640 (2004).

26. L. Gattinoni et al., Wnt signaling arrests effector T cell differentiation and generates CD8+ memory stem cells. Nat Med 15, 808–813 (2009).

27. C. Yao et al., BACH2 enforces the transcriptional and epigenetic programs of stem-like CD8. Nat Immunol 22, 370–380 (2021).

28. I. H. Sun et al., mTOR Complex 1 Signaling Regulates the Generation and Function of Central and Effector Foxp3. J Immunol 201, 481–492 (2018).

29. N. Hayatsu et al., Analyses of a Mutant Foxp3 Allele Reveal BATF as a Critical Transcription Factor in the Differentiation and Accumulation of Tissue Regulatory T Cells. Immunity 47, 268–283.e269 (2017).

30. C. Neumann et al., c-Maf-dependent T. Nat Immunol 20, 471–481 (2019).

31. A. J. Lam, P. Uday, J. K. Gillies, M. K. Levings, Helios is a marker, not a driver, of human Treg stability. Eur J Immunol 52, 75–84 (2022).

32. M. A. Burchill, J. Yang, C. Vogtenhuber, B. R. Blazar, M. A. Farrar, IL-2 receptor beta-dependent STAT5 activation is required for the development of Foxp3+ regulatory T cells. J Immunol 178, 280–290 (2007).

33. G. J. Martinez et al., The transcription factor NFAT promotes exhaustion of activated CD8⁺ T cells. Immunity 42, 265–278 (2015).

34. G. P. Mognol et al., Exhaustion-associated regulatory regions in CD8. Proc Natl Acad Sci U S A 114, E2776–E2785 (2017).

35. H. Seo et al., TOX and TOX2 transcription factors cooperate with NR4A transcription factors to impose CD8. Proc Natl Acad Sci U S A 116, 12410–12415 (2019).

36. A. G. Levine, A. Arvey, W. Jin, A. Y. Rudensky, Continuous requirement for the TCR in regulatory T cell function. Nat Immunol 15, 1070–1078 (2014).

37. T. Chinen et al., An essential role for the IL-2 receptor in T. Nat Immunol 17, 1322–1333 (2016).

38. S. Fisson et al., Continuous activation of autoreactive CD4+ CD25+ regulatory T cells in the steady state. J Exp Med 198, 737–746 (2003).

39. C. L. Tan et al., PD-1 restraint of regulatory T cell suppressive activity is critical for immune tolerance. J Exp Med 218, (2021).

40. G. Wang, M. Tajima, T. Honjo, A. Ohta, STAT5 interferes with PD-1 transcriptional activation and affects CD8+ T-cell sensitivity to PD-1-dependent immunoregulation. Int Immunol 33, 563–572 (2021).

41. K. H. Toomer et al., Essential and non-overlapping IL-2Rα-dependent processes for thymic development and peripheral homeostasis of regulatory T cells. Nat Commun 10, 1037 (2019).

42. J. A. Perry et al., PD-L1-PD-1 interactions limit effector regulatory T cell populations at homeostasis and during infection. Nat Immunol 23, 743–756 (2022).

43. O. S. Qureshi et al., Constitutive clathrin-mediated endocytosis of CTLA-4 persists during T cell activation. J Biol Chem 287, 9429–9440 (2012).

44. H. Schneider, C. E. Rudd, Diverse mechanisms regulate the surface expression of immunotherapeutic target ctla-4. Front Immunol 5, 619 (2014).

45. N. K. Serwas et al., Human DEF6 deficiency underlies an immunodeficiency syndrome with systemic autoimmunity and aberrant CTLA-4 homeostasis. Nat Commun 10, 3106 (2019).

46. D. Janman et al., Regulation of CTLA-4 recycling by LRBA and Rab11. Immunology 164, 106–119 (2021).

47. K. Hildner et al., Batf3 deficiency reveals a critical role for CD8alpha+ dendritic cells in cytotoxic T cell immunity. Science 322, 1097–1100 (2008).

48. J. C. Cancel, K. Crozat, M. Dalod, R. Mattiuz, Are Conventional Type 1 Dendritic Cells Critical for Protective Antitumor Immunity and How? Front Immunol 10, 9 (2019).

49. T. L. Murphy, K. M. Murphy, Dendritic cells in cancer immunology. Cell Mol Immunol 19, 3–13 (2022).

50. F. Mattei, G. Schiavoni, F. Belardelli, D. F. Tough, IL-15 is expressed by dendritic cells in response to type I IFN, double-stranded RNA, or lipopolysaccharide and promotes dendritic cell activation. J Immunol 167, 1179–1187 (2001).

51. J. Liu, X. Zhang, Y. Cheng, X. Cao, Dendritic cell migration in inflammation and immunity. Cell Mol Immunol 18, 2461–2471 (2021).

52. K. Rawat et al., CCL5-producing migratory dendritic cells guide CCR5+ monocytes into the draining lymph nodes. J Exp Med 220, (2023).

53. C. Oderup, L. Cederbom, A. Makowska, C. M. Cilio, F. Ivars, Cytotoxic T lymphocyte antigen-4-dependent down-modulation of costimulatory molecules on dendritic cells in CD4+ CD25+ regulatory T-cell-mediated suppression. Immunology 118, 240–249 (2006).

54. Y. Onishi, Z. Fehervari, T. Yamaguchi, S. Sakaguchi, Foxp3+ natural regulatory T cells preferentially form aggregates on dendritic cells in vitro and actively inhibit their maturation. Proc Natl Acad Sci U S A 105, 10113–10118 (2008).

55. W. Kastenmuller et al., Regulatory T cells selectively control CD8+ T cell effector pool size via IL-2 restriction. J Immunol 187, 3186–3197 (2011).

56. M. Tekguc, J. B. Wing, M. Osaki, J. Long, S. Sakaguchi, Treg-expressed CTLA-4 depletes CD80/CD86 by trogocytosis, releasing free PD-L1 on antigen-presenting cells. Proc Natl Acad Sci U S A 118, (2021).

57. F. Marangoni et al., Expansion of tumor-associated Treg cells upon disruption of a CTLA-4-dependent feedback loop. Cell 184, 3998–4015.e3919 (2021).

58. S. D. Spencer et al., The orphan receptor CRF2-4 is an essential subunit of the interleukin 10 receptor. J Exp Med 187, 571–578 (1998).

59. D. J. Berg et al., Interleukin-10 is a central regulator of the response to LPS in murine models of endotoxic shock and the Shwartzman reaction but not endotoxin tolerance. J Clin Invest 96, 2339–2347 (1995).

60. A. De Gassart et al., MHC class II stabilization at the surface of human dendritic cells is the result of maturation-dependent MARCH I down-regulation. Proc Natl Acad Sci U S A 105, 3491–3496 (2008).

61. L. E. Tze et al., CD83 increases MHC II and CD86 on dendritic cells by opposing IL-10-driven MARCH1-mediated ubiquitination and degradation. J Exp Med 208, 149–165 (2011).

62. S. T. Ferris et al., A minor subset of Batf3-dependent antigen-presenting cells in islets of Langerhans is essential for the development of autoimmune diabetes. Immunity 41, 657–669 (2014).

63. J. A. Carrero et al., Resident macrophages of pancreatic islets have a seminal role in the initiation of autoimmune diabetes of NOD mice. Proc Natl Acad Sci U S A 114, E10418–E10427 (2017).

64. P. N. Zakharov, H. Hu, X. Wan, E. R. Unanue, Single-cell RNA sequencing of murine islets shows high cellular complexity at all stages of autoimmune diabetes. J Exp Med 217, (2020).

65. L. A. Kalekar et al., CD4(+) T cell anergy prevents autoimmunity and generates regulatory T cell precursors. Nat Immunol 17, 304–314 (2016).

66. K. Pletinckx et al., Immature dendritic cells convert anergic nonregulatory T cells into Foxp3-IL-10+ regulatory T cells by engaging CD28 and CTLA-4. Eur J Immunol 45, 480–491 (2015).

67. A. S. Thomann et al., Conversion of Anergic T Cells Into Foxp3. Front Immunol 12, 704578 (2021).

68. D. M. Gravano, D. A. Vignali, The battle against immunopathology: infectious tolerance mediated by regulatory T cells. Cell Mol Life Sci 69, 1997–2008 (2012).

69. R. Setoguchi, S. Hori, T. Takahashi, S. Sakaguchi, Homeostatic maintenance of natural Foxp3(+) CD25(+) CD4(+) regulatory T cells by interleukin (IL)-2 and induction of autoimmune disease by IL-2 neutralization. J Exp Med 201, 723–735 (2005).

70. E. E. West et al., PD-L1 blockade synergizes with IL-2 therapy in reinvigorating exhausted T cells. J Clin Invest 123, 2604–2615 (2013).

71. S. Ma et al., Attenuated IL-2 muteins leverage the TCR signal to enhance regulatory T cell homeostasis and response. Front Immunol 14, 1257652 (2023).

72. T. Duhen, R. Duhen, A. Lanzavecchia, F. Sallusto, D. J. Campbell, Functionally distinct subsets of human FOXP3+ Treg cells that phenotypically mirror effector Th cells. Blood 119, 4430–4440 (2012).

73. J. B. Wing, A. Tanaka, S. Sakaguchi, Human FOXP3. Immunity 50, 302–316 (2019).

74. M. L. Alegre et al., Regulation of surface and intracellular expression of CTLA4 on mouse T cells. J Immunol 157, 4762–4770 (1996).

75. X. B. Wang, C. Y. Zheng, R. Giscombe, A. K. Lefvert, Regulation of surface and intracellular expression of CTLA-4 on human peripheral T cells. Scand J Immunol 54, 453–458 (2001).

76. E. T. Hayes, C. E. Hagan, L. Khoryati, M. A. Gavin, D. J. Campbell, Regulatory T Cells Maintain Selective Access to IL-2 and Immune Homeostasis despite Substantially Reduced CD25 Function. J Immunol 205, 2667–2678 (2020).

77. K. W. Hwang et al., Transgenic expression of CTLA-4 controls lymphoproliferation in IL-2-deficient mice. J Immunol 173, 5415–5424 (2004).

78. M. E. Raeber, R. A. Rosalia, D. Schmid, U. Karakus, O. Boyman, Interleukin-2 signals converge in a lymphoid-dendritic cell pathway that promotes anticancer immunity. Sci Transl Med 12, (2020).

79. S. C. Eisenbarth, Dendritic cell subsets in T cell programming: location dictates function. Nat Rev Immunol 19, 89–103 (2019).

80. A. J. Schottelius, M. W. Mayo, R. B. Sartor, A. S. Baldwin, Interleukin-10 signaling blocks inhibitor of kappaB kinase activity and nuclear factor kappaB DNA binding. J Biol Chem 274, 31868–31874 (1999).

81. M. Zagorulya et al., Tissue-specific abundance of interferon-gamma drives regulatory T cells to restrain DC1-mediated priming of cytotoxic T cells against lung cancer. Immunity 56, 386–405.e310 (2023).

82. M. A. Moreno Ayala et al., CXCR3 expression in regulatory T cells drives interactions with type I dendritic cells in tumors to restrict CD8. Immunity 56, 1613–1630.e1615 (2023).

83. M. S. Anderson, J. A. Bluestone, The NOD mouse: a model of immune dysregulation. Annu Rev Immunol 23, 447–485 (2005).

84. Y. P. Lai et al., CD28 engagement inhibits CD73-mediated regulatory activity of CD8. Commun Biol 4, 595 (2021).

85. M. Martínez-Llordella et al., CD28-inducible transcription factor DEC1 is required for efficient autoreactive CD4+ T cell response. J Exp Med 210, 1603–1619 (2013).

86. G. Cheng et al., IL-2 receptor signaling is essential for the development of Klrg1+ terminally differentiated T regulatory cells. J Immunol 189, 1780–1791 (2012).

87. J. Yan, B. Liu, Y. Shi, H. Qi, Class II MHC-independent suppressive adhesion of dendritic cells by regulatory T cells in vivo. J Exp Med 214, 319–326 (2017).

88. A. Butler, P. Hoffman, P. Smibert, E. Papalexi, R. Satija, Integrating single-cell transcriptomic data across different conditions, technologies, and species. Nat Biotechnol 36, 411–420 (2018).

89. M. E. Ritchie et al., limma powers differential expression analyses for RNA-sequencing and microarray studies. Nucleic Acids Res 43, e47 (2015).

90. D. Wu et al., ROAST: rotation gene set tests for complex microarray experiments. Bioinformatics 26, 2176–2182 (2010).

91. M. D. Robinson, A. Oshlack, A scaling normalization method for differential expression analysis of RNA-seq data. Genome Biol 11, R25 (2010).

