## Supplemental Figures for "An IL-2 mutein increases IL-10 and CTLA-4-dependent suppression of dendritic cells by regulatory T cells"

**FIGURE S1**

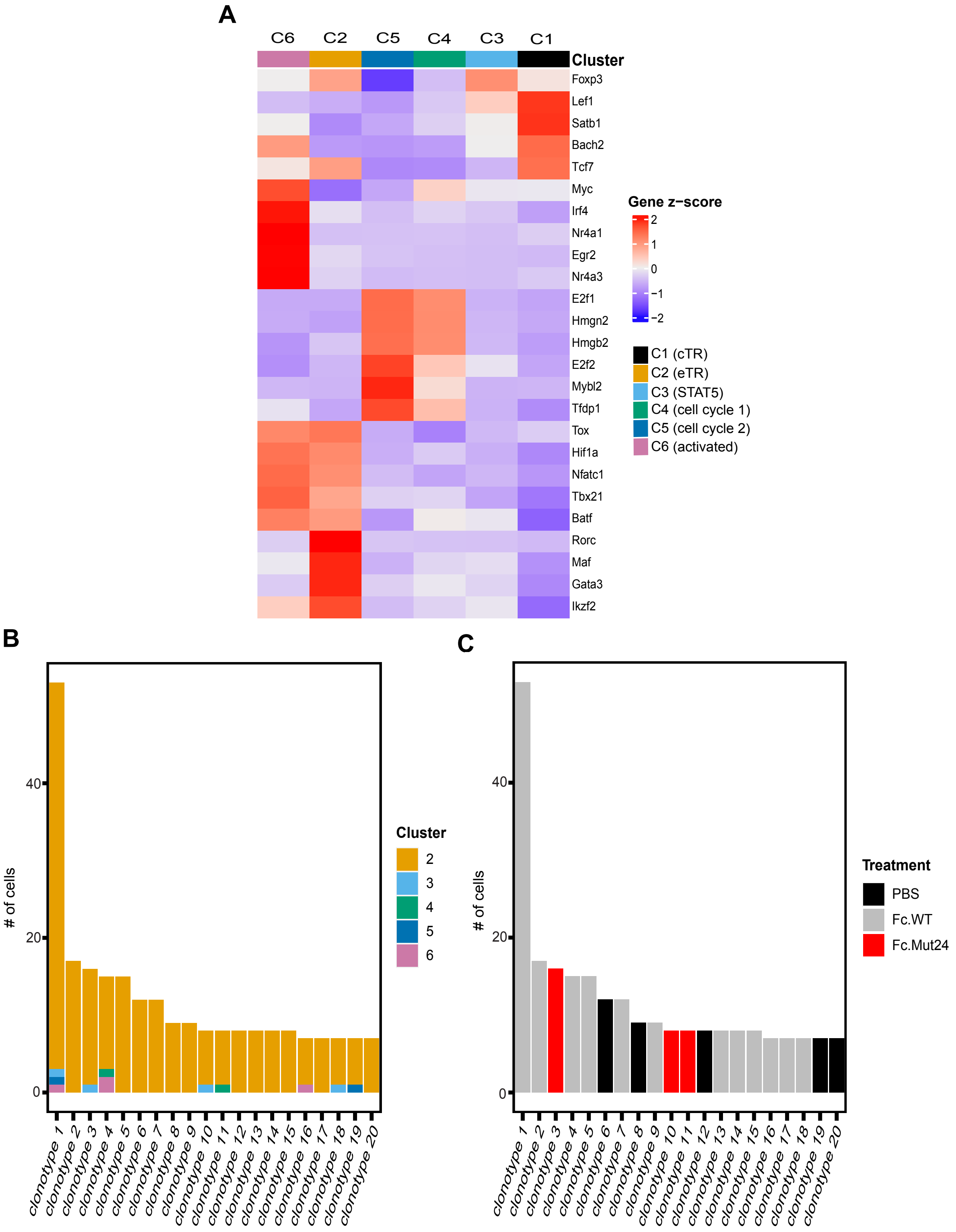


**Figure S1. Fc.Mut24 does not impact Foxp3$^+$ T$_R$ cell clonal expansion, related to Figure 1.** Splenocytes from C57BL/6 Foxp3-mRFP treated with PBS, Fc.WT, or Fc.Mut24 were harvested 3 days later and sorted to obtain CD4$^+$ Foxp3$^+$ regulatory T (T$_R$) cells for single-cell RNA-seq (scRNA-seq). **A.** Heat map showing the mean expression (z-score) of representative transcription factor genes (rows) for each cluster (columns). **B&C.** Bar graphs showing the number of cells identified for each of the top twenty T$_R$ clonotypes colored by cluster **(B)** or treatment group **(C).**

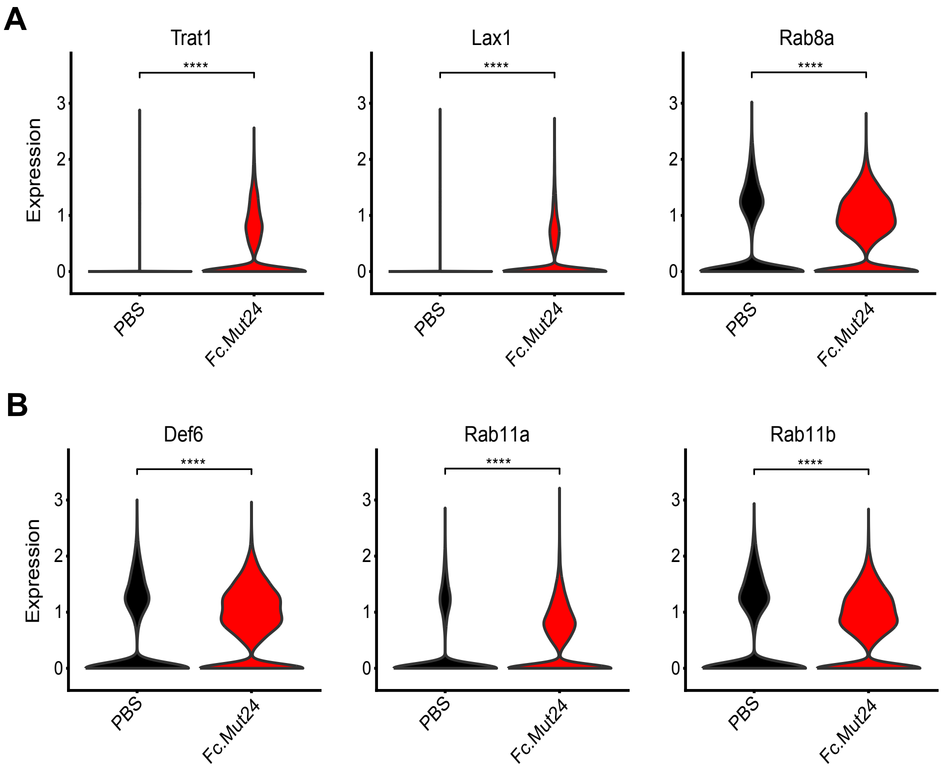


**Figure S2. Fc.Mut24 increases the expression of genes involved in CTLA-4 trafficking and recycling in Foxp3⁺ $T_R$ cells, related to Figure 1 and 3.** Splenocytes from C57BL/6 Foxp3-mRFP treated with PBS, Fc.WT, or Fc.Mut24 were harvested 3 days later and sorted to obtain $CD4^+$ $Foxp3^+$ regulatory T ($T_R$) cells for single-cell RNA-seq (scRNA-seq). Violin plots showing normalized expression for the indicated genes involved in CTLA-4 trafficking to the cell surface **(A)** or endosomal recycling of CTLA-4 **(B)**.

**FIGURE S3**

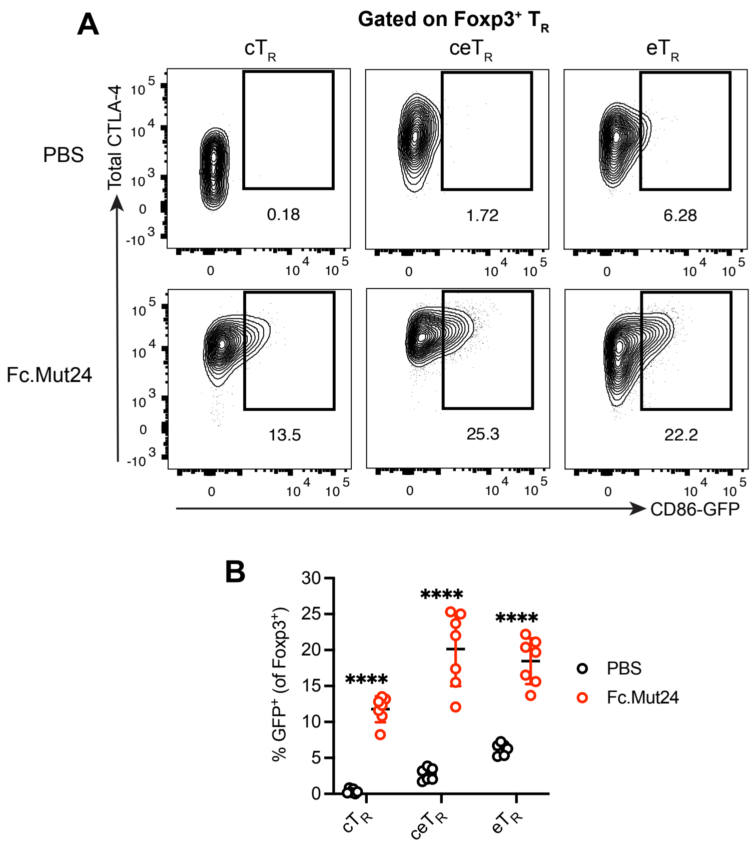


**Figure S3. Fc.Mut24 increases CTLA-4-dependent transendocytosis by different Foxp3⁺ $T_R$ cell populations, related to Figure 4. A.** Capture of CD86-GFP by CD44$^{lo}$ CD62L$^{hi}$ central $T_R$ (c$T_R$), CD44$^{hi}$ CD62L$^{hi}$ central effector $T_R$ (ce$T_R$), and CD44$^{hi}$ CD62L$^{lo}$ effector $T_R$ (e$T_R$) cells between different treatment groups. See Figure 4A for the experimental design schematic. Representative flow cytometry plots are shown. **B.** Graph shows mean ± SD with individual data points (n = 7); ****$P \leq 0.0001$, multiple unpaired *t*-tests. Data are representative of 2-3 independent experiments.

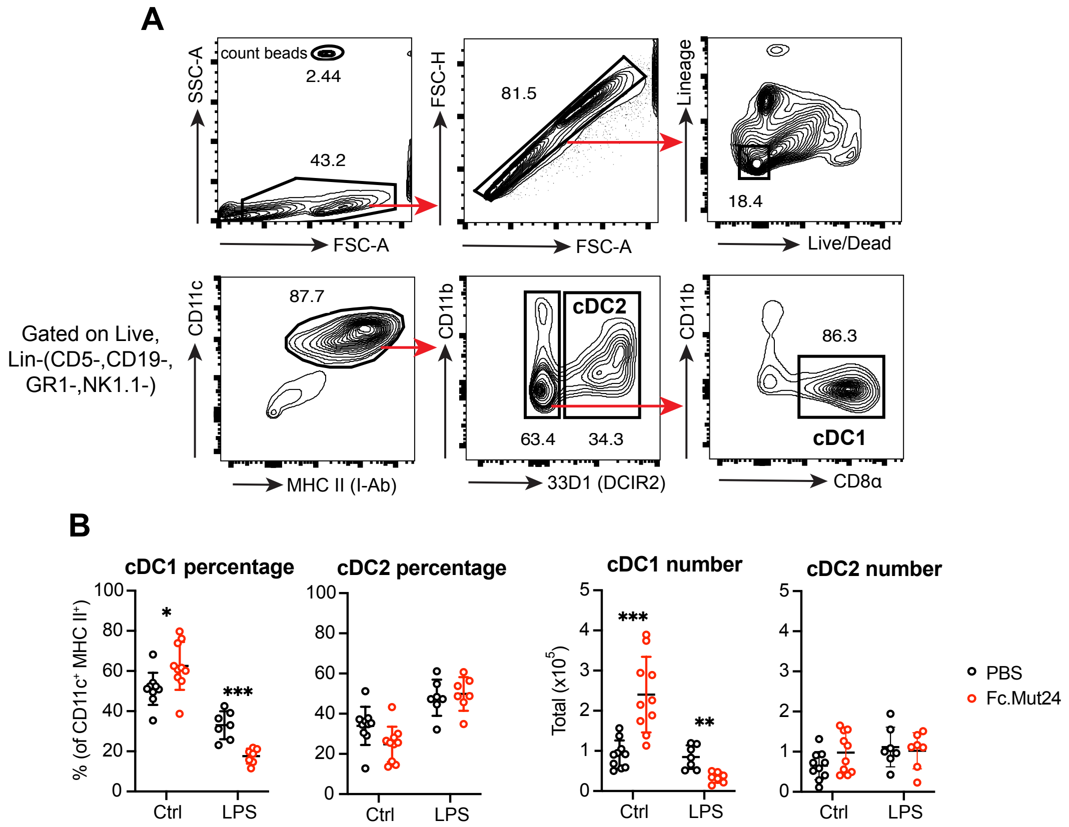


**Figure S4. Fc.Mut24 expands cDC1s in C57BL/6 mice, related to Figure 5. A.** Representative gating strategy for CD11c$^+$MHC II$^+$33D1$^-$ CD8$\alpha^+$ cDC1 and CD11c$^+$MHC II$^+$33D1$^+$ cDC2. **B.** Percentage and number of cDC1 and cDC2 between different treatment groups. See Figure 5A for the experimental design schematic. Graphs show mean ± SD with individual data points (n = 7 to 10); *$P \leq 0.05$, **$P \leq 0.01$, ***$P \leq 0.001$, multiple unpaired *t*-tests. Ctrl, Control. Data are representative of 2-3 independent experiments.

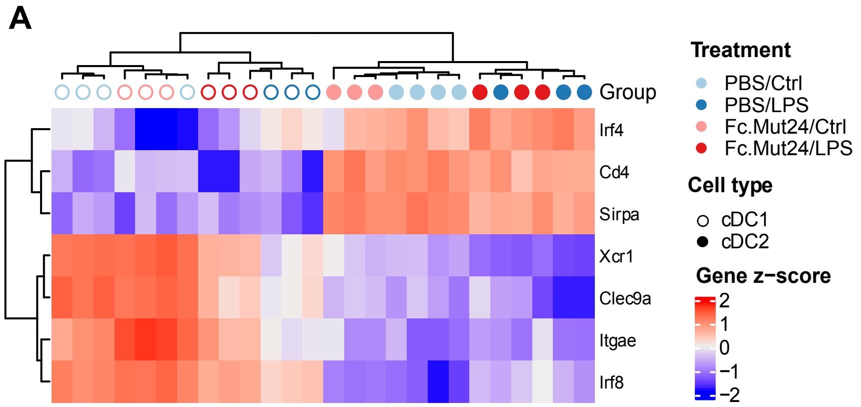


**Figure S5. Expression of known identity markers in dendritic cells after Fc.Mut24 treatment, related to Figure 5.** Unstimulated (Ctrl) or LPS-stimulated splenocytes from C57BL/6 mice treated with PBS or Fc.Mut24 were harvested 4 days post-treatment and sorted to obtain cDC1 or cDC2 populations for bulk RNA-seq. See Figure 5A for the experimental design schematic**. A.** Heatmap showing the mean expression (z-score) of select genes differentially expressed between cDC1 and cDC2 for all treatment groups.

FIGURE S6

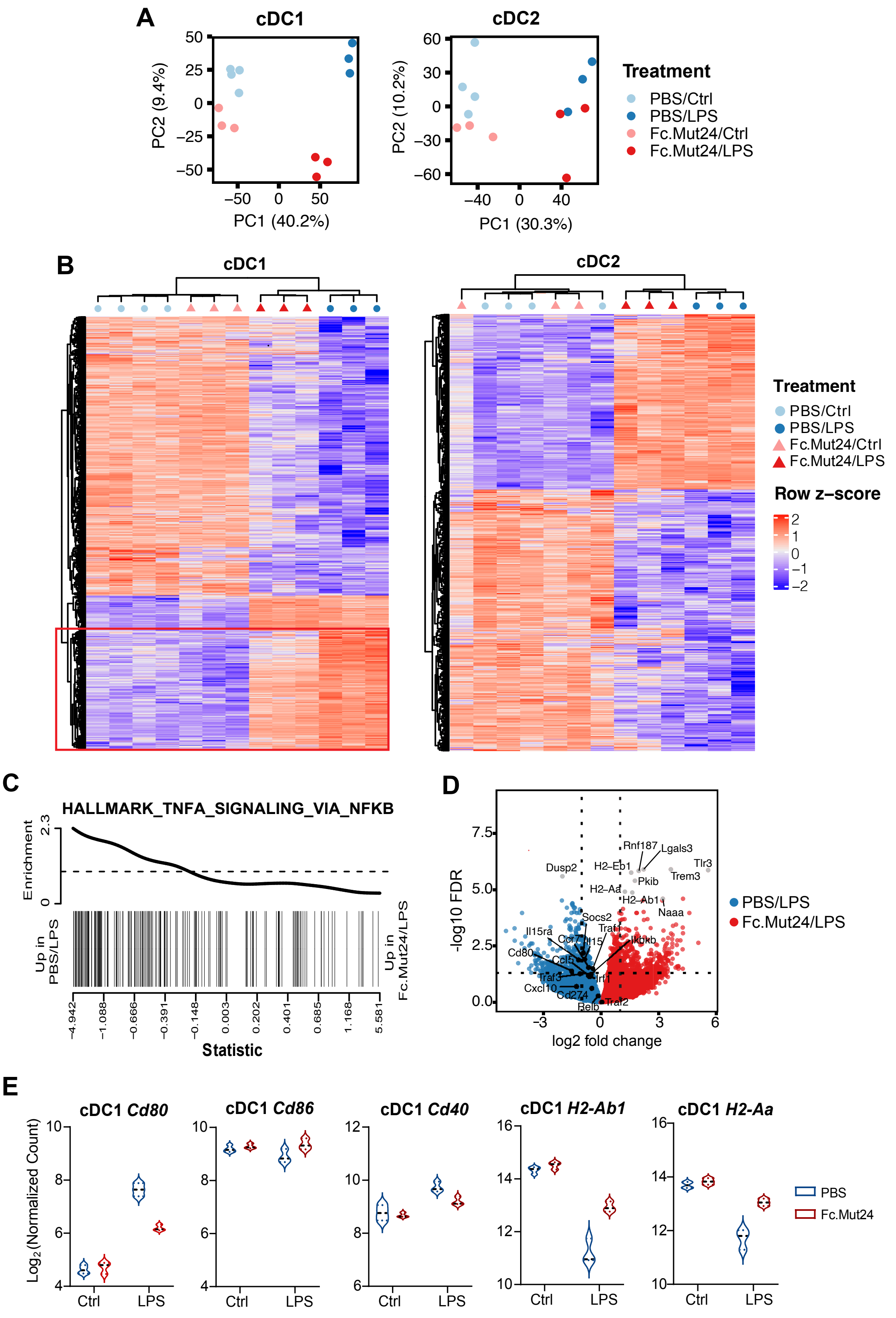
A
cDC1
cDC2
PC2 (9.4%)
PC1 (40.2%)
PC2 (10.2%)
PC1 (30.3%)
Treatment
PBS/Ctrl
PBS/LPS
Fc.Mut24/Ctrl
Fc.Mut24/LPS
B
cDC1
cDC2
Treatment
PBS/Ctrl
PBS/LPS
Fc.Mut24/Ctrl
Fc.Mut24/LPS
Row z–score
C
HALLMARK_TNFA_SIGNALING_VIA_NFKB
Enrichment
Up in PBS/LPS
Up in Fc.Mut24/LPS
Statistic
D
-log10 FDR
log2 fold change
PBS/LPS
Fc.Mut24/LPS
E
cDC1 Cd80
cDC1 Cd86
cDC1 Cd40
cDC1 H2-Ab1
cDC1 H2-Aa
Log2(Normalized Count)
Ctrl
LPS
PBS
Fc.Mut24

**Figure S6. Impact of Fc.Mut24 treatment of the LPS-mediated transcriptional response in dendritic cells, related to Figure 5.** Unstimulated (Ctrl) or LPS-stimulated splenocytes from C57BL/6 mice treated with PBS or Fc.Mut24 were harvested 4 days post-treatment and sorted to obtain cDC1 or cDC2 populations for bulk RNA-seq. See Figure 5A for the experimental design schematic**. A.** Principal component analysis of global transcriptional signatures from indicated cell types and treatment groups. **B.** Heatmap showing the mean expression (z-score) of all genes differentially expressed between LPS and Ctrl samples in cDC1 and cDC2 (FDR < 0.01, $Log_2$ fold-change > 1). In cDC1s, a subset of genes (red rectangle) have diminished LPS-mediated upregulation in the presence of Fc.Mut24. **C.** Gene set enrichment analysis of cDC1s showing that the Hallmark TNFA Signaling via NFKB pathway is enriched in PBS/LPS relative to Fc.Mut24/LPS samples (FDR = 0.005). **D.** Volcano plot of cDC1s comparing Fc.Mut24 treatment versus PBS with the top ten differentially expressed genes by FDR (gray) and select NFκB-regulated genes (black) highlighted. **E.** Violin plots of cDC1s showing $Log_2$ normalized counts for select genes: *Cd80*, *Cd86*, *CD40*, *H2-Ab1* (I-$A^b$ beta chain), and *H2-Aa* (I-$A^b$ alpha chain).

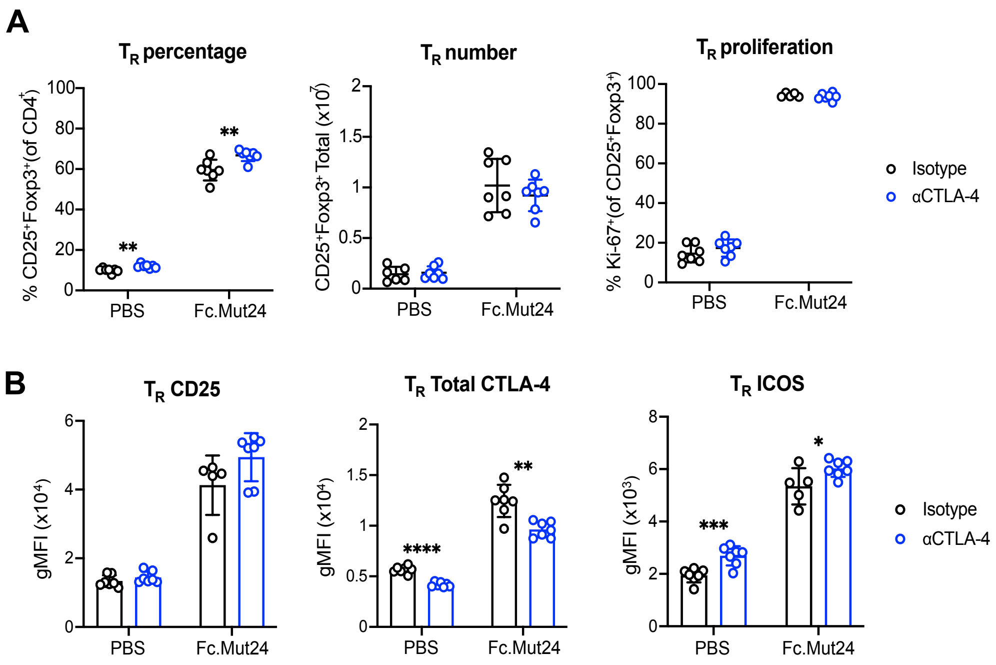


**Figure S7. The anti-CTLA-4 antibody 4F10 does not cause Foxp3$^+$ $T_R$ cell depletion during Fc.Mut24 treatment, related to Figure 5.** Splenocytes from C57BL/6 mice treated with PBS or Fc.Mut24 followed by isotype or anti-CTLA-4 antibody and LPS stimulation were analyzed by flow cytometry. Gates were set on live, CD4$^+$ cells. **A&B.** Graphs show mean ± SD with individual data points (n = 7); **P* ≤ 0.05, ***P* ≤ 0.01, ****P* ≤ 0.001, *****P* ≤ 0.0001, multiple unpaired *t*-tests. Data are representative of 2-3 independent experiments. **B.** Gates were set on CD25$^+$ Foxp3$^+$ $T_R$ for analysis of the expression of indicated $T_R$ activation markers.

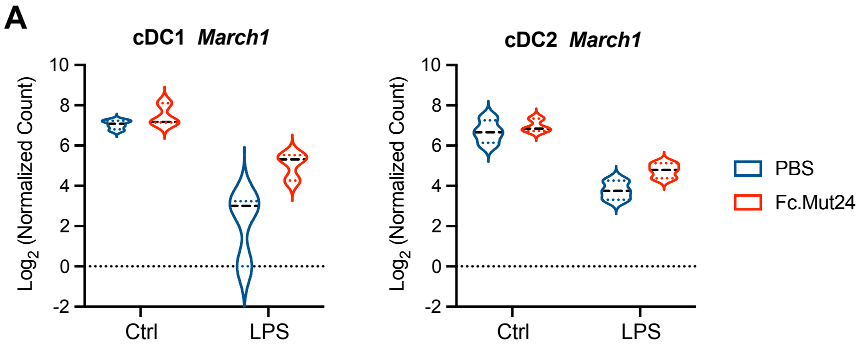


**Figure S8. Fc.Mut24 increases March1 gene expression in cDC1s from C57BL/6 mice, related to Figure 6.** Unstimulated (Ctrl) or LPS-stimulated splenocytes from C57BL/6 mice treated with PBS or Fc.Mut24 were harvested 4 days post-treatment and sorted to obtain cDC1 or cDC2 populations for bulk RNA-seq. See Figure 5A for the experimental design schematic **A.** Violin plots showing $Log_2$ normalized counts for *March1* gene expression between treatment groups.

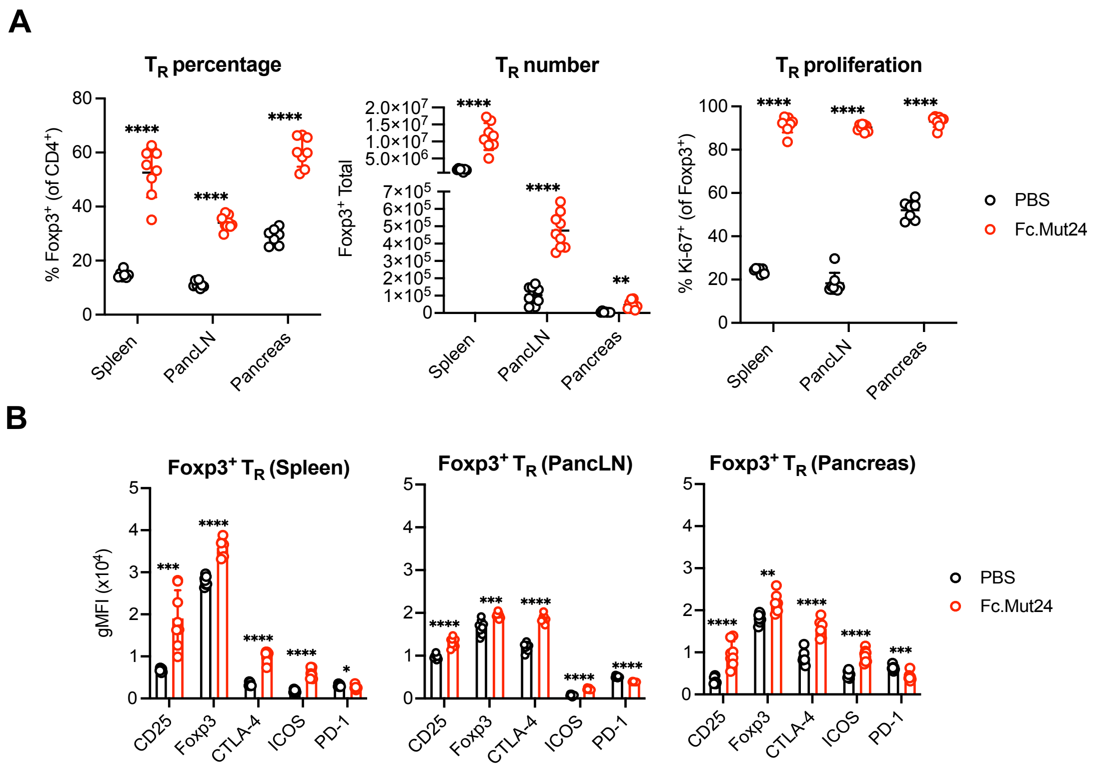


**Figure S9. Fc.Mut24 promotes Foxp3⁺ $T_R$ cell activation in NOD mice, related to Figure 7.** The spleen, pancreatic lymph node (PancLN), and pancreas of 8-10-week-old female non-obese diabetic (NOD) mice treated with PBS or Fc.Mut24 were harvested 4 days later and analyzed by flow cytometry. Gates were set on live, CD45⁺, CD4⁺ cells. Percentage, number, and proliferation (Ki-67⁺) of Foxp3⁺ $T_R$ cells **(A)** and expression of indicated $T_R$ activation markers **(B)**. **A&B.** All graphs show mean ± SD with individual data points (n = 6 to 9); **P* ≤ 0.05, ***P* ≤ 0.01, ****P* ≤ 0.001, *****P* ≤ 0.0001, multiple unpaired *t*-tests. Data are representative of 2-3 independent experiments.

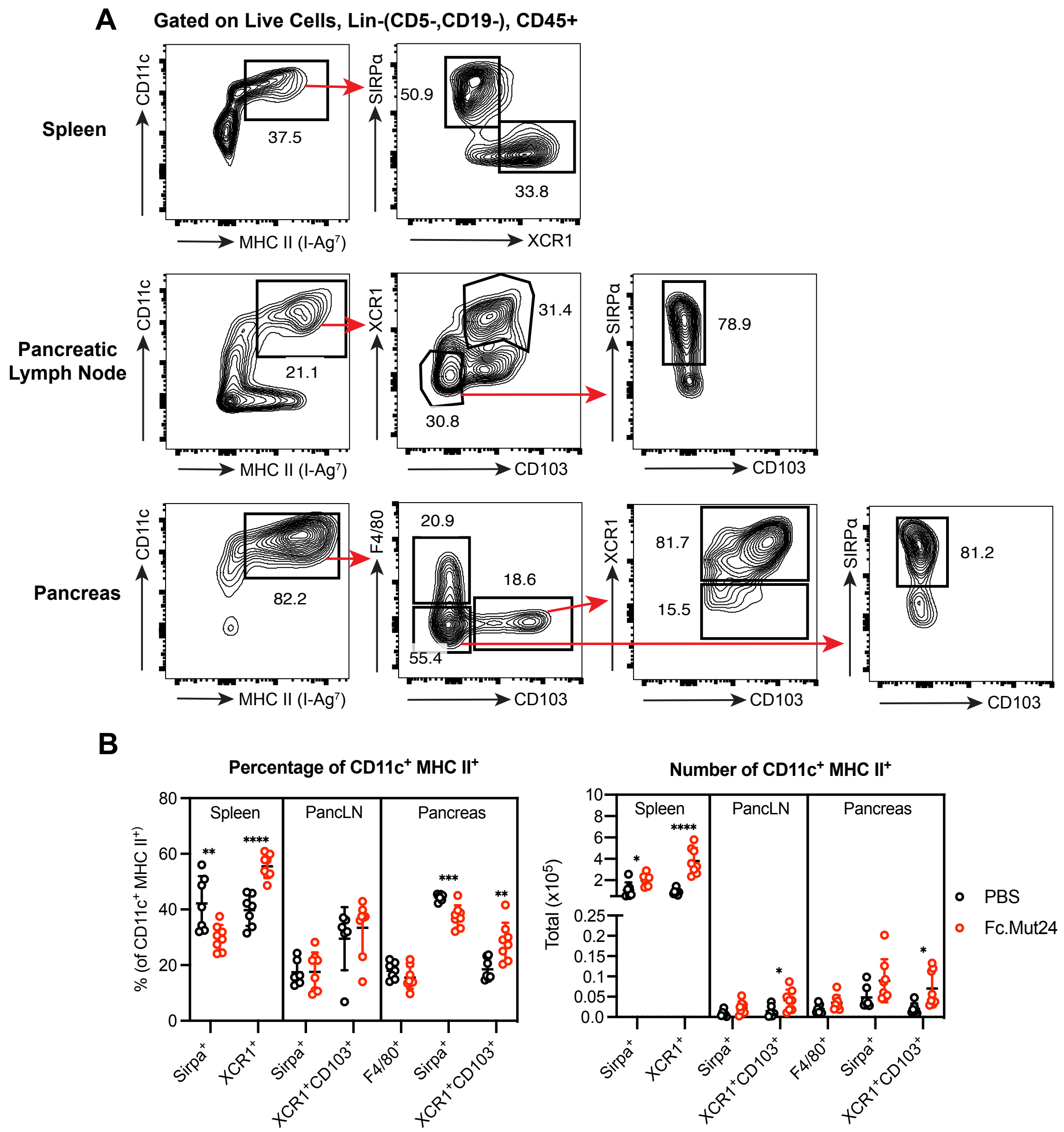


**Figure S10. Fc.Mut24 expands cDC1s in NOD mice, related to Figure 7.** The spleen, pancreas, and pancreatic lymph node (PancLN) of 8-10-week-old female non-obese diabetic (NOD) mice treated with PBS or Fc.Mut24 were harvested 4 days later and analyzed by flow cytometry. **A.** Representative gating strategy for the CD11c$^+$ MHC II$^+$ antigen-presenting cell subsets in different tissues. **B.** Percentage and number of different subsets of CD11c$^+$ MHC II$^+$ antigen-presenting cells. Graphs show mean ± SD with individual data points (n = 6 to 8); *$P \leq 0.05$, **$P \leq 0.01$, ***$P \leq 0.001$, ****$P \leq 0.0001$, multiple unpaired *t*-tests. Data are representative of 2-3 independent experiments.

**Table S1. Full list of DEGs between each Foxp3$^+$ $T_R$ cell cluster from the scRNA-seq experiment, related to Figure 1B&C.** See Table S1.xlsx.
